# Signals in the Cells: Multimodal and Contextualized Machine Learning Foundations for Therapeutics

**DOI:** 10.1101/2024.06.12.598655

**Authors:** Alejandro Velez-Arce, Xiang Lin, Michelle M. Li, Kexin Huang, Wenhao Gao, Tianfan Fu, Bradley L. Pentelute, Manolis Kellis, Marinka Zitnik

**Affiliations:** Department of Biomedical Informatics, Harvard Medical School, Boston, MA 02115; Department of Computer Science, Stanford School of Engineering, Stanford, CA 94305; Department of Chemical Engineering, Massachusetts Institute of Technology, Cambridge, MA 02139; Department of Computational Science, Rensselaer Polytechnic Institute, Troy, NY 12180; Department of Chemistry, Massachusetts Institute of Technology, Cambridge, MA 02139; Broad Institute of MIT and Harvard, Computer Science and Artificial Intelligence Laboratory, MIT, Electrical Engineering and Computer Science Department, Massachusetts Institute of Technology, Cambridge, MA 02139; Broad Institute of MIT and Harvard, Harvard Data Science Initiative, Kempner Institute for the Study of Natural and Artificial Intelligence, Harvard University, Department of Biomedical Informatics, Harvard Medical School, Boston, MA 02215

## Abstract

Drug discovery AI datasets and benchmarks have not traditionally included single-cell analysis biomarkers. While benchmarking efforts in single-cell analysis have recently released collections of single-cell tasks, they have yet to comprehensively release datasets, models, and benchmarks that integrate a broad range of therapeutic discovery tasks with cell-type-specific biomarkers. Therapeutics Commons (TDC-2) presents datasets, tools, models, and benchmarks integrating cell-type-specific contextual features with ML tasks across therapeutics. We present four tasks for contextual learning at single-cell resolution: drug-target nomination, genetic perturbation response prediction, chemical perturbation response prediction, and protein-peptide interaction prediction. We introduce datasets, models, and benchmarks for these four tasks. Finally, we detail the advancements and challenges in machine learning and biology that drove the implementation of TDC-2 and how they are reflected in its architecture, datasets and benchmarks, and foundation model tooling.

## 1 Introduction

Single-cell genomics has enabled the study of cellular processes with remarkable resolution, offering insights into cellular heterogeneity and dynamics [1]. Progress in data generation and computational methods designed for single-cell analysis [2] has facilitated machine learning models that incorporate cell-type-specific data across various therapeutic areas [3, 4]. Despite these advances, there remains a need for comprehensive datasets, benchmarks, and tools that integrate single-cell analysis with diverse therapeutic approaches.

Out-of-distribution (OOD) generalization [5] and incorporation of novel tools [6] and modalities [4, 7] remain as challenges for biomedical machine learning models across a broad range of tasks. Models capable of accurate OOD predictions promise to expand to the vast molecular space, whose size is estimated at 10^60^ potential drug-like molecules [8], yet less than 10^5^ of those are FDA-approved drugs [9], suggesting the potential for advanced computational methods to navigate the molecular space and help find, generate, and optimize candidate drugs. Further, handling multimodal data is essential for building foundation models that accurately capture the complex interactions within biological systems [10], which is vital for understanding disease mechanisms and discovering effective treatments. These challenges are compounded by the lack of unified datasets organized across stages of drug discovery. Therapeutics Data Commons [11, 12] addresses these challenges by providing a unified platform that consolidates therapeutic datasets and benchmarks. Still, benchmarks tailored to measuring the effectiveness of models at OOD predictions are rare for several key biological tasks [13]. Most dataset and benchmark providers also struggle to evaluate models using longitudinal data [14] and real-world evidence [15] due to challenges in continual data collection [16].

Public benchmarks and competitions that measure performance using state-of-the-art methods against standardized criteria have a strong track record of accelerating innovation in algorithm development in therapeutic science [17]. However, machine learning researchers face several challenges in this domain, including: (1) a lack of domain knowledge regarding essential tasks in the field, (2) the absence of standard benchmarks for different methods due to their varying implementations [17], and (3) the high cost of implementing complex data pre-processing pipelines for each task [11]. Despite the progress made over the last five years in developing datasets and benchmarks for machine learning methods in therapeutics [18, 12], decentralized and standardized benchmarks are still needed in single-cell therapeutics. Similarly, recent advancements in single-cell analysis tools, datasets, and benchmarks [1, 2, 19] have yet to address therapeutic tasks.

Further, there is demand for retrieval tooling that can integrate into emerging tool-based LLMs [20, 21]. [21] shows that integrating retrieval APIs with LLMs mitigates the issue of hallucination, commonly encountered when prompting LLMs directly. They also discuss the challenges of supporting a web scale collection of millions of changing APIs. With the advent of public petabyte-scale public-API-accessible databases in biomedical informatics [2], it is imperative to build systems providing unified retrieval across biomedical modalities and large-scale, changing, data sources.

### Present work

The Commons 2.0, a.k.a. TDC-2, introduces a multi-modal retrieval API implementing a novel API-first-dataset architecture. The API-first-dataset architecture supports retrieval of continually updated heterogenous data sources (i.e., knowledge graphs [22], embedding models [19], petabyte-scale RNA data [2], etc.) across biomedical modalities providing abstractions for complex data processing pipelines [11], data lineage [23], and versioning [24]. This lays the foundation for improving the stability of biomedical AI workflows with continuous data updates [25]. We have developed these resources building on the Therapeutic Data Commons platform, and leverage them to present four novel ML tasks with fine-grained biological contexts: single-cell drug-target identification [4], single-cell chemical/genetic perturbation response predictions [26, 27], and a cell-type-specific protein-peptide interaction task [28, 15] (Section 3.3.1). The models, datasets, and benchmarks composing these tasks address cell-type-specific molecule ML modeling [4], evaluation of contextual AI models [29, 4], heuristics for generating negative samples in peptidomimetics [30], and OOD generalization in single-cell perturbation response prediction [31, 32]. Overall, the contributions are (summarized in Table 5):

1. TDC-2 is the first platform to integrate single-cell analysis with multimodal machine learning in drug discovery via four contextual AI tasks along with corresponding models, datasets, and benchmarks (Section 3).
2. TDC-2 formalizes contextualized metrics for evaluating contextual AI models on therapeutic tasks. This allows for evaluating models’ abilities in identifying the most predictive cell type contexts (Section 4).
3. TDC-2 introduces an API-first-dataset architecture (section 6.3.1) for augmenting therapeutic models with retrieval APIs, improving workflow stability with continual data collection.
4. An unprecedented collection of heterogeneous data sources is unified under the API-first-dataset architecture (section 6.3.2). These include: a petabyte-scale single-cell RNA data atlas [2], a framework for biomedical knowledge graphs [22], and a collection of retrieval APIs integrated with complex data processing workflows (Section 6.3.2; see listing 2 for example usage).

## 2 Related Work

### Machine learning datasets and benchmarks in therapeutics

Therapeutics Data Commons (TDC) was the first unifying platform providing systematic access and evaluation for machine learning across the entire range of therapeutics [11]. TDC included 66 AI-ready datasets and 22 learning tasks, spanning the discovery and development of safe and effective medicines. TDC also provided an ecosystem of tools and community resources, including 33 data functions and types of meaningful data splits, 23 strategies for systematic model evaluation, 17 molecule generation oracles, and 29 public leaderboards. TDC-2 augments the biomedical modalities covered by TDC data, tasks, and benchmarks to lay the foundations for building and evaluating foundation models. We expanded the biomedical modalities covered by TDC, introduced single-cell resolution to various modalities, introduced access to model embeddings, introduced machine learning model retrieval APIs for inference and fine-tuning, and incorporated contextualized metrics into model evaluation to lay the foundations for building and evaluating single-cell therapeutic foundation models. We further developed an API-first dataset design (section 6.3.1) unifying modalities [33] for augmenting LLMs with biomedical experimental data retrieval [6]. TDC-2 distinguishes itself from related datasets [34, 35], benchmarks [17, 18, 36, 37], model development frameworks [38, 39, 40], and therapeutic initiatives [2] in its integration of single-cell analysis with multimodal machine learning in drug discovery via four contextual AI tasks and retrieval APIs for multimodal datasets and models.

### Therapeutic and single-cell foundation models

Foundation models trained on TDC, and a subset of tasks in TDC-2 (Sections 7.2.3, 7.2.4, and 7.2.5), have been shown to generalize across several therapeutic tasks [41]. Additionally, parallel efforts in training foundation models on large single-cell atlases have shown a potential to advance cell type annotation and matching of healthy-disease cells to study cellular signatures of disease [3, 42, 43, 44]. TDC-2 bridges these independent and parallel efforts by providing formal definitions of therapeutic tasks, datasets, and benchmarks at single-cell resolution in order to provide precise therapeutic predictions incorporating cell type contexts [29, 4].

### Augmenting biomedical AI workflows with multi-modal tooling

Recent advancements in tool-based large language models (LLMs) showcase the potential of augmenting these systems to call external functions and APIs [20, 21]. By integrating chemistry tooling, LLMs can be augmented with novel capabilities across chemical tasks [6]. LLM multi-agent frameworks have also succeeded at manipulating collections of tools for the automatic processing and execution of single-cell analysis tasks [45]. The API-first [46, 47, 48] approach adopted by TDC-2’s multimodal retrieval API, dubbed the API-first dataset, integrates expert-designed tools with continually updated data to support grounding of biomedical AI workflows.

## 3 Results

TDC-2 introduces four tasks with fine-grained biological contexts: contextualized drug-target identification, single-cell chemical/genetic perturbation response prediction, and cell-type-specific protein-peptide binding interaction prediction, which introduce antigen-processing-pathway-specific, cell-type-specific, peptide-specific, and patient-specific biological contexts. Benchmarks for drug target nomination, genetic perturbation response prediction and chemical perturbation response prediction, all at single-cell resolution, were computed with corresponding leaderboards introduced on the TDC website. In addition, a benchmark and leaderboard was introduced for the TCR-Epitope binding interaction task for peptide design at single-cell resolution for T-cell receptors.

### Context-specific metrics

In real-world machine learning applications, data subsets can correspond to critical outcomes. In therapeutics, there is evidence that the effects of drugs can vary depending on the type of cell they are targeting and where specific proteins are acting [49]. We build on the “slice” abstraction [50] to measure model performance at critical biological subsets. Context-specific metrics are defined to measure model performance at critical biological slices, with our benchmarks focused on measuring cell-type-specific model performance. In the case of benchmarks for perturbation response prediction and protein-peptide binding affinity, the studies are limited to a particular cell line, however our definition for context-specific metrics lays the foundations for building models which can generalize across cell lines and make context-aware predictions [29, 50]. For single-cell drug-target nomination, we measure model performance at top-performing cell types. See Section 7.2.6 for definitions.

### 3.2 TDC.scDTI: Contextualized Drug-Target Identification

#### Motivation

Single-cell data have enabled the study of gene expression and function at the level of individual cells across healthy and disease states [51, 2, 29]. To facilitate biological discoveries using single-cell data, machine-learning models have been developed to capture the complex, cell-type-specific behavior of genes [3, 44, 42, 4]. In addition to providing the single-cell measurements and foundation models, TDC-2 supports the development of contextual AI models to nominate therapeutic targets in a cell type-specific manner [4]. We introduce a benchmark dataset, model, and leaderboard for context-specific therapeutic target prioritization, encouraging the innovation of model architectures (e.g., to incorporate new modalities, such as protein structure and sequences [52, 53, 54, 55, 56], genetic perturbation data [57, 58, 59, 60], disease-specific single-cell atlases [61, 62, 63], and protein networks [64, 65, 66]). TDC-2’s release of TDC.scDTI is a step in standardizing benchmarks for more comprehensive assessments of context-specific model performance.

### 3.2 Dataset and benchmark

We use curated therapeutic target labels from the Open Targets Platform [67] for rheumatoid arthritis (RA) and inflammatory bowel disease (IBD) [4] (section 7.1.1). We benchmark PINNACLE [4]—trained on cell type specific protein-protein interaction networks—and a graph attention neural network (GAT) [68]—trained on a context-free reference protein-protein interaction network—on the curated therapeutic targets dataset. As expected, PINNACLE underperforms when evaluated on context-agnostic metrics (Table 1) and drastically outperforms GAT when evaluated on context-specific metrics (Appendix Table 1). To our knowledge, TDC-2 provides the first benchmark for context-specific learning [29]. TDC-2’s contribution helps standardize the evaluation of single-cell ML models for drug target identification and other single-cell tasks [42, 3, 4, 44].

**Figure 1:**
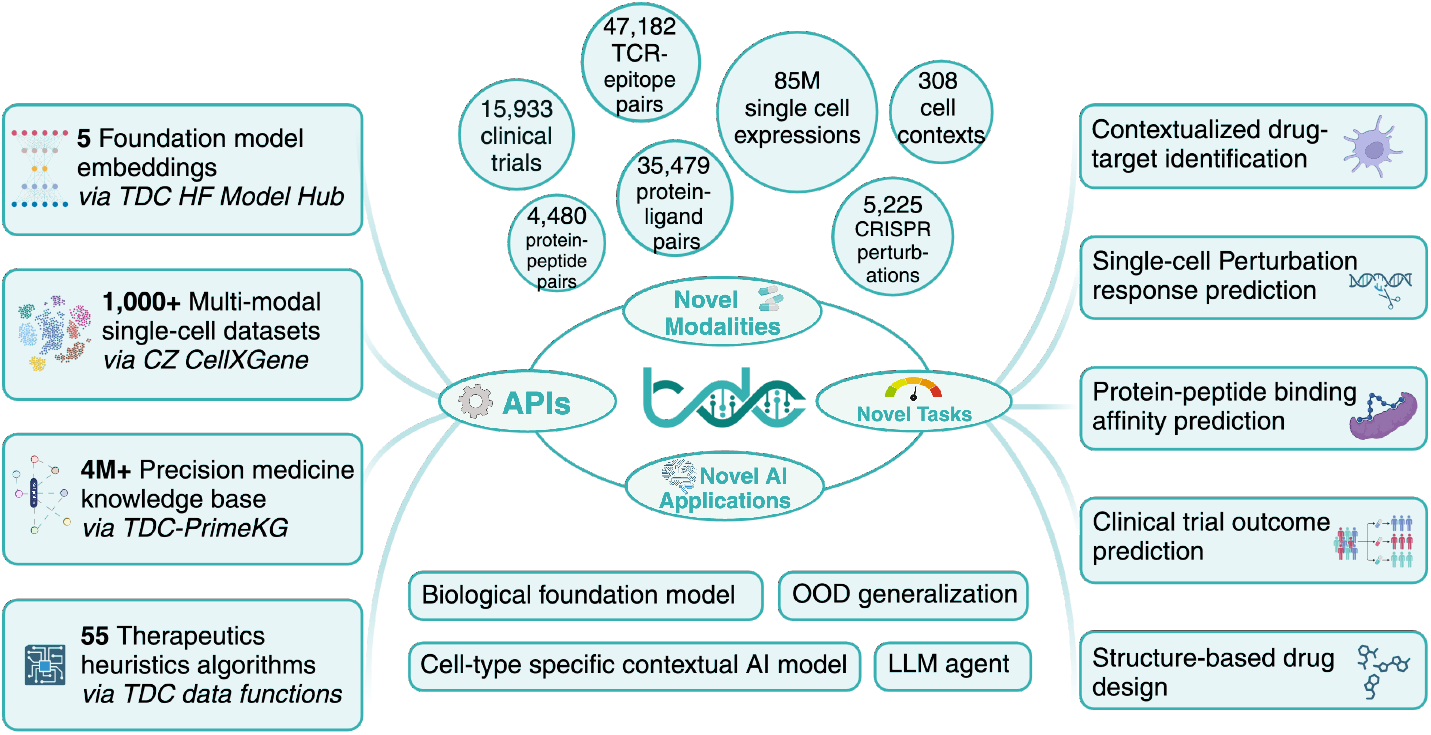
Overview of TDC-2. TDC-2 introduces a multimodal retrieval API powering ML-task-driven [11] datasets [69, 4, 67, 107, 91, 90, 14, 15, 109, 108]and benchmarks spanning 10+ new modalities and 5 state-of-the-art machine learning tasks (section 7.2), including 4 contextual AI tasks: TDC.scDTI (section 3.1), single-cell genetic perturbation response prediction (section 3.2.1), single-cell chemical perturbation response prediction (section 3.2.2), and single-cell protein-peptide interaction prediction (section 3.3). Model benchmarks highlighting biomedical AI challenges in OOD Generalization [26, 27, 120, 14] and evaluation [4, 30] of cell-type-specific contextual AI models are introduced.

**Table 1:** Cell-type specific target nomination for two therapeutic areas, rheumatoid arthritis (RA) and inflammatory bowel diseases (IBD). Cell-type specific context metrics (definitions in Section 7.2.6): AP@5 Top-20 CT -average precision at k = 5 for the 20 best-performing cell types (CT); AUROC Top-1 CT - AUROC for top-performing cell type; AUROC Top-10 CT and AUROC Top-20 CT - weighted average AUROC for top-10 and top-20 performing cell types, respectively, each weighted by the number of samples in each cell type; AP@5/AUROC CF - context-free AP@5/AUROC integrated across all cell types. Shown are results from models run on ten independent seeds. N/A - not applicable.

| Model | AP@5 Top-20 CT | AUROC Top-1 CT | AUROC Top-10 CT | AUROC Top-20 CT | AP@5 CF | AUROC CF |
| --- | --- | --- | --- | --- | --- | --- |
| PINNACLE (RA) | 0.913±0.059 | 0.765±0.054 | 0.676±0.017 | 0.647±0.014 | 0.226±0.023 | 0.510±0.005 |
| GAT (RA) | N/A | N/A | N/A | N/A | 0.220±0.013 | 0.580±0.010 |
| PINNACLE (IBD) | 0.873±0.069 | 0.935±0.067 | 0.799±0.017 | 0.752±0.011 | 0.198±0.013 | 0.500±0.010 |
| GAT (IBD) | N/A | N/A | N/A | N/A | 0.200±0.023 | 0.640±0.017 |

### 3.2 TDC.PerturbOutcome: Perturbation-Response Prediction

#### Motivation

Understanding and predicting transcriptional responses to genetic or chemical pertur-bations provides insights into cellular adaptation and response mechanisms. Such predictions can advance therapeutic strategies, as they enable researchers to anticipate how cells will react to targeted interventions, potentially guiding more effective treatments. Models that have shown promise at this task [26, 27] are limited to either genetic or chemical perturbations without being able to generalize to the other. Approaches that can generalize across chemical and genetic perturbations [31, 32] may be unable to generalize to unseen perturbations without modification.

#### Dataset and benchmark

We used the scPerturb [69] datasets to benchmark the generalizability of perturbation-response prediction models across seen/unseen perturbations and cell lines. We benchmark models in genetic and chemical perturbations using metrics measuring intra/inter-cell line and seen/unseen perturbation generalizability. We provide results measuring unseen perturbation generalizability for Gene Perturbation Response Prediction using the scPerturb gene datasets (Norman K562, Replogle K562, Replogle RPE1) [70, 71] with results shown in Table 2. For Chemical Perturbation Prediction, we evaluated chemCPA utilizing cold splits on perturbation type and showed a significant decrease in performance for 3 of 4 perturbations evaluated (3). We have also included Biolord [31] and scGen [72] for comprehensive benchmarking on the well-explored perturbation response prediction on seen perturbation types problem. These tests were run on sciPlex2 [73].

**Table 2:** Unseen genetic perturbation response prediction. We evaluate GEARS across the top 20 differentially expressed genes, based on the highest absolute differential expression upon perturbation, for MSE (MSE@20DEG). Gene expression was measured in log normalized counts. In single-cell analysis, a standard procedure is to normalize the counts within each cell so that they sum to a specific value (usually the median sum across all cells in the dataset) and then to log transform the values using the natural logarithm [26]. For both normalization and ranking genes by differential expression, we utilized Scanpy [74]. We used the sc.tl.rank_genes_groups() function with default parameters in Scanpy, which employs a t-test to estimate scores. This function provides a z-score for each gene and ranks genes based on the absolute values of the score. Genes showing a significant level of dropout were not included in this metric.

| Dataset | Tissue | Cell Line | Method | MSE@20DEG |
| --- | --- | --- | --- | --- |
| Norman K562 | K562 | lymphoblast | no-perturb | 0.341±0.001 |
| Norman K562 | K562 | lymphoblast | CPA | 0.230±0.008 |
| Norman K562 | K562 | lymphoblast | GEARS | 0.176±0.003 |
| Replogle 562 | K562 | lymphoblast | no-perturb | 0.126±0.000 |
| Replogle 562 | K562 | lymphoblast | CPA | 0.126±0.000 |
| Replogle 562 | K562 | lymphoblast | GEARS | 0.109±0.004 |
| Replogle RPE1 | RPE-1 | epithelial | no-perturb | 0.164±0.000 |
| Replogle RPE1 | RPE-1 | epithelial | CPA | 0.162±0.001 |
| Replogle RPE1 | RPE-1 | epithelial | GEARS | 0.110±0.003 |

##### 3.2.1 Genetic Perturbation Response Prediction

We use scPerturb gene datasets (Norman K562, Replogle K562, Replogle RPE1) [70, 71]. In the case of single-gene perturbations, we assessed the models based on the perturbation of experimentally perturbed genes not included in the training data. We used data from two genetic perturbation screens, with 1,543 perturbations for RPE-1 (retinal pigment epithelium) cells and 1,092 for K-562 cells, each involving over 170,000 cells. These screens utilized the Perturb-seq assay, which combines pooled screening with single-cell RNA sequencing to analyze the entire transcriptome for each cell. We trained GEARS separately on each dataset. In addition to an existing deep learning-based model (CPA), we also developed a baseline model (no perturbation), assuming that gene expression remains unchanged after perturbation.

We evaluated the models’ performance by calculating the mean squared error between the predicted gene expression after perturbation and the actual post-perturbation expression for the held-out set. Based on the highest absolute differential expression upon perturbation, the top 20 most differentially expressed genes were selected.

##### 3.2.2 Chemical Perturbation Response Prediction

The dataset consists of four drug-based perturbations from sciPlex2 [73, 69] (BMS, Dex, Nutlin, SAHA). sciPlex2 contains alveolar basal epithelial cells from the A549 (lung adenocarcinoma), K562 (chronic myelogenous leukemia), and MCF7 (mammary adenocarcinoma) tissues. Results are shown in Table 3. Our experiments rely on the coefficient of determination (R^2^) as the primary performance measure. We calculate this score by comparing actual measurements with counterfactual predictions for all genes. Assessing all genes is essential to evaluating the decoder’s overall performance and understanding the background context. Still, it is also beneficial to determine performance based on top differentially expressed genes [27]. The baseline used discards all perturbation information, adequately measuring the improvement resulting by the models’ drug encoding [27]. ChemCPA’s performance dropped by an average of 15% across the four perturbations. The maximum drop was 34%. Code for intra/inter cell-line benchmarks for chemical (drug) and genetic (CRISPR) perturbations is in Appendix 7.3.5 and Appendix 7.3.5, respectively. Using this code, users can evaluate models of their choice on the benchmark and submit them to the TDC-2 leaderboards for this task (Appendix 7.3.5).

**Table 3:** Unseen chemical perturbation response prediction. We have evaluated chemCPA utilizing cold splits on perturbation type and show a significant decrease in performance for 3 of 4 perturbations evaluated. We have also included Biolord [31] and scGen [72] for comparison. The dataset used consists of four chemical (drug) perturbations from sciPlex2 [69] (BMS, Dex, Nutlin, SAHA). sciPlex2 contains alveolar basal epithelial cells from the A549 (lung adenocarcinoma), K562 (chronic myelogenous leukemia), and MCF7 (mammary adenocarcinoma) tissues. Our experiments rely on the coefficient of determination (R^2^) as the primary performance measure.

| Drug | Method | $R^2$ (seen perturbations) | $R^2$ (unseen perturbations) |
| --- | --- | --- | --- |
| BMS | Baseline | 0.620±0.044 | N/A |
| Dex | Baseline | 0.603±0.053 | N/A |
| Nutlin | Baseline | 0.628±0.036 | N/A |
| SAHA | Baseline | 0.617±0.027 | N/A |
| BMS | Biolord | 0.939±0.022 | N/A |
| Dex | Biolord | 0.942±0.028 | N/A |
| Nutlin | Biolord | 0.928±0.026 | N/A |
| SAHA | Biolord | 0.980±0.005 | N/A |
| BMS | ChemCPA | 0.943±0.006 | 0.906±0.006 |
| Dex | ChemCPA | 0.882±0.014 | 0.540±0.013 |
| Nutlin | ChemCPA | 0.925±0.010 | 0.835±0.009 |
| SAHA | ChemCPA | 0.825±0.026 | 0.690±0.021 |
| BMS | scGen | 0.903±0.030 | N/A |
| Dex | scGen | 0.944±0.018 | N/A |
| Nutlin | scGen | 0.891±0.032 | N/A |
| SAHA | scGen | 0.948±0.034 | N/A |

### 3.3 TDC.ProteinPeptide: Contextualized Protein-Peptide Interaction Prediction

#### Motivation

Evaluating protein-peptide binding prediction models requires standardized benchmarks, presenting challenges in assessing and validating model performance across different studies [13]. Despite the availability of several benchmarks for protein-protein interactions, this is not the case for protein-peptide interactions. The renowned multi-task benchmark for Protein sEquence undERstanding (PEER) [18] and MoleculeNet [35] both lack support for a protein-peptide interaction prediction task. Furthermore, protein-peptide binding mechanisms vary wildly by cellular and biological context [75, 76, 77, 78]. Current models, as such, tend to be restricted to one task instance (i.e., T Cell Receptor (TCR) and Peptide-MHC Complex or B Cell Receptor (BCR) and Antigen Peptide binding) and do not span protein-peptide interactions [79, 80, 81, 7, 82, 83, 84].

We define and evaluate a subtask for TCR-Epitope binding interaction prediction applying contextual AI (Section 7.2.3) to the T Cell cell line.

#### 3.3.1 TCR-Epitope (Peptide-MHC Complex) Interaction Prediction

The critical challenge in TCR-Epitope (Peptide-MHC Complex) interaction prediction lies in creating a model that can effectively generalize to unseen TCRs and epitopes [85]. While TCR-H [86] and TEINet [87] have shown improved performance on prediction for known epitopes, by incorporating advanced features like attention mechanisms and transfer learning, the performance considerably drops for unseen epitopes [88, 89]. Another challenge in TCR-Epitope interaction prediction lies in the choice of heuristic for generating negative samples, with non-binders often underrepresented or biased in curated datasets, leading to inaccurate predictions when generalized [30].

### Datasets and Benchmarks

TDC-2 establishes a curated dataset and benchmark within its single-cell protein-peptide binding affinity prediction task to measure model generalizability to unseen TCRs and epitopes and model sensitivity to the selection of negative data points. Benchmarking datasets use three types of heuristics for generating negative samples: random shuffling of epitope and TCR sequences (RN), experimental negatives (NA), and pairing external TCR sequences with epitope sequences (ET). We harness data from the TC-hard dataset [90] for the first two types and PanPep [91] for the third type. Both datasets use hard [90] splits, ensuring that epitopes in the testing set are not present in the training set. Our results (Table 4) show the lack of a reasonable heuristic for generating negative samples, with model performance evaluation shown to be unsatisfactory. For two heuristics, all models perform poorly. The best-performing model in ET is MIX-TPI, with roughly 0.70 AUROC. The best-performing model in RN is AVIB-TCR, with approximately 0.576 AUROC. For NA, 4 of 6 models perform near-perfectly as measured on AUROC. Models benchmarked include AVIB-TCR [28], MIX-TPI [92], Net-TCR2 [93], PanPep [85], TEINet [87], and TITAN [89].

**Table 4:**
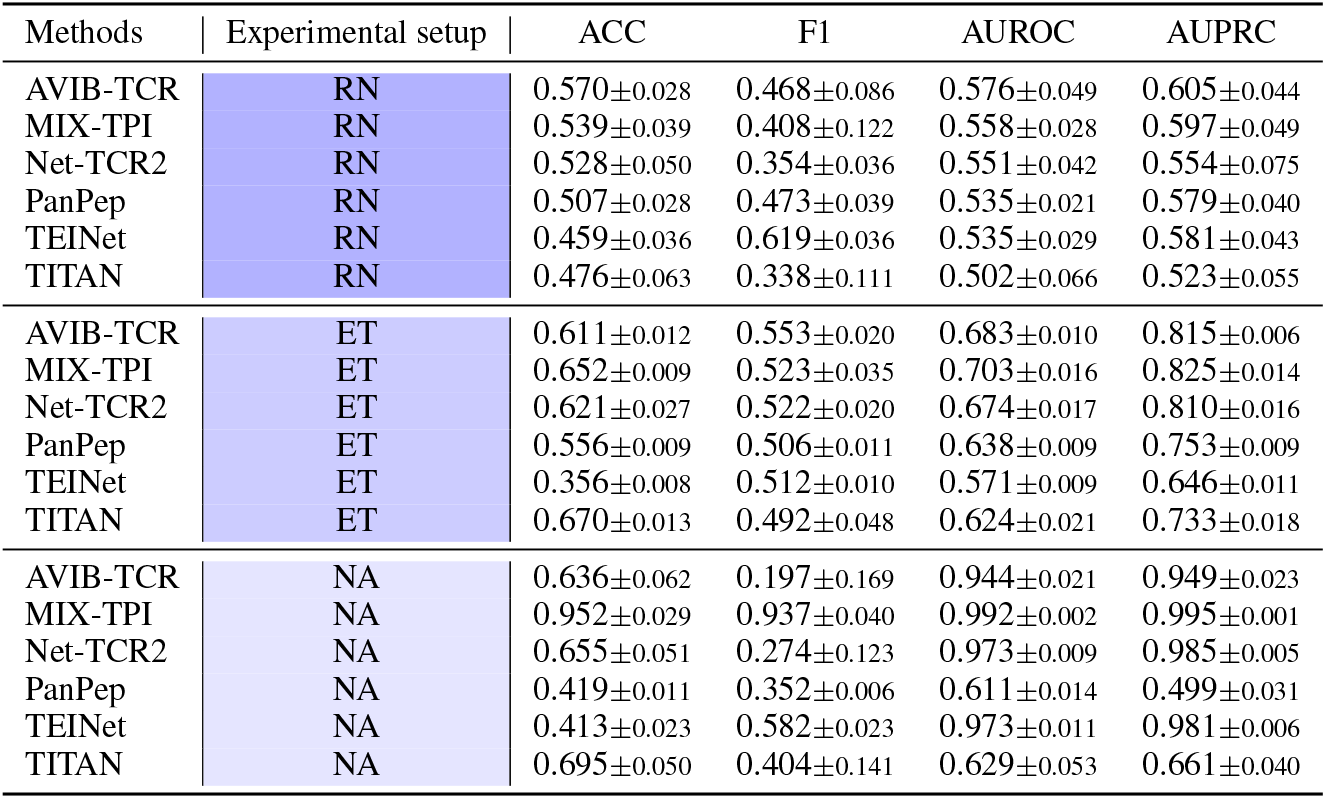
TCR-epitope binding interaction binary classification performance. All models perform poorly under realistic but challenging RN and ET experimental setups. The best-performing model in RN is AVIB-TCR, with an average of 0.576 (AUROC). The best-performing model in ET is MIX-TPI, with an average of 0.700 (AUROC). For NA, 4 of 6 models achieve near-perfect AUROC.

**Table 5:** Comparison of TDC-2 with other datasets, benchmarks, and ML platforms in therapeutics TDC-2 distinguishes itself from related datasets [34, 35], benchmarks [17, 18, 36, 37], model development frameworks [38, 39, 40], and therapeutic initiatives [2] in its integration of single-cell analysis with multimodal machine learning in drug discovery via four contextual AI tasks and retrieval APIs for multimodal datasets and models. The API-first [46, 47, 48] approach adopted by TDC-2’s multimodal retrieval API, dubbed the API-first dataset, enables development of large language models invoking experimental biomedical data retrieval APIs based on therapeutic queries. TDC-2 integrates expert-designed tools into this unified LLM-friendly API to foster scientific advancement by bridging the gap between experimental and computational therapeutic science. OP - Open Problems.

| Feature | TDC | TorchDrug | MoleculeNet | OP | scPerturb | CELLXGENE | TDC-2 |
| --- | --- | --- | --- | --- | --- | --- | --- |
| Single-cell Drug-Target Identification Task | X | X | X | X | X | X | ✓ |
| CRISPR-Cell Perturbation Response Prediction | X | X | X | X | ✓ | X | ✓ |
| Drug-Cell Perturbation Response Prediction | X | X | X | X | ✓ | X | ✓ |
| Single-cell Protein-Peptide Binding Interaction Prediction | X | X | X | X | X | X | ✓ |
| Structure-Based Drug Design | X | ✓ | ✓ | X | X | X | ✓ |
| Clinical Trial Outcome Prediction | X | X | X | X | X | X | ✓ |
| Knowledge Graphs | X | ✓ | X | X | X | X | ✓ |
| Python API | ✓ | ✓ | X | ✓ | X | ✓ | ✓ |
| Single-cell Gene Expression Atlases | X | X | X | ✓ | ✓ | ✓ | ✓ |
| External API data retrieval | X | X | X | X | X | X | ✓ |
| Data Views | X | X | X | X | X | X | ✓ |
| Foundation Model Embedding Retrieval | X | ✓ | X | X | X | ✓ | ✓ |

**Table 6:** Clinical Trial Outcome Prediction task benchmark model results on the TOP dataset [14], described in section 6.4.1.

| Model | Phase 1 AUPRC | Phase 2 AUPRC | Phase 3 AUPRC | Indication Level AUPRC |
| --- | --- | --- | --- | --- |
| HINT | 0.772 | 0.607 | 0.623 | 0.703 |
| COMPOSE | 0.665 | 0.532 | 0.545 | 0.624 |
| DeepEnroll | 0.701 | 0.580 | 0.590 | 0.655 |

## 4 Discussion

TDC-2 makes technical strides in unifying a challenging body of work under a representative architecture (section 6.3.1) and a collection of datasets (Section 7.1), benchmarks (Section 3), and foundation model tools (Section 6.3.2). The presented models and benchmarks illustrate the challenges of developing machine learning methods in therapeutics capable of generalizing to out-of-distribution samples across varying biological contexts.

### Identifying most predictive biological contexts and cell types

There is evidence that the effects of drugs can vary depending on the type of cell they are targeting and where specific proteins are acting [49]. Therefore, it is essential to identify the most predictive cell-type contexts by evaluating therapeutic machine-learning tasks with cell-type-specific metrics. This approach can help determine the cell types that play crucial and distinct roles in the disease pathogenesis of conditions like rheumatoid arthritis (RA) and inflammatory bowel diseases (IBD). In our study, we used cell-type-specific metrics to compare the performance of [4] and [68] models for the TDC.scDTI task (Table 1). The results showed that PINNACLE protein representations outperformed the GAT model for RA and IBD diseases across the top 1, 10, and 20 (most predictive) cell types.

### Generalizing genetic perturbation response prediction models to out-of-distribution samples

Gene editing is a promising tool for diseases that cannot be effectively treated or managed with alternative therapeutic modalities. For instance, the FDA recently approved gene editing to modify T-cells for treating patients with acute lymphoblastic leukemia [94]. However, many disease-causing genetic variants result from insertions and deletions, making it crucial to accurately predict gene editing outcomes to ensure effectiveness and minimize off-target effects. Understanding how cells respond to genetic changes is essential for this. However, the vast number of potential genetic changes makes it difficult to study all possibilities experimentally [95]. Although there have been improvements in predicting the outcomes of genetic changes, applying these predictions to new genetic changes and cell types when predicting gene expression responses remains challenging [26]. We assessed the performance of various models in predicting gene responses to single and multiple genetic perturbations using single-cell RNA-sequencing data from genetic screens. We found that recently developed models perform well when tested on a single cell line; however, they struggle to generalize to cell lines not encountered during training. This gap presents valuable opportunities for algorithmic innovation.

### Generalizing chemical perturbation response prediction models to out-of-distribution samples

Using high-throughput techniques such as cell hashing has made it easier to conduct single-cell RNA sequencing in multi-sample experiments at a low cost. However, these methods require expensive library preparation and are not easily scalable to many perturbations. This becomes incredibly challenging when studying the effects of combination therapies or genetic perturbations, where experimental screening of all possible combinations becomes impractical. While projects like the Human Cell Atlas aim to comprehensively map cellular states across tissues, creating a similar atlas for the effects of perturbations on gene expression is impossible due to the vast number of possibilities. Therefore, it is crucial to develop computational tools to guide the exploration of the perturbation space and identify promising candidate combination therapies in high-throughput screenings. A successful computational method for navigating the perturbation space should be able to predict cell behavior when subject to novel combinations of perturbations that were only measured separately in the original experiment. In our study, we benchmarked Biolord [31], scGen [72], and ChemCPA [27] on chemical perturbation response prediction and showed significant improvement over the baseline method. Furthermore, we demonstrate a considerable drop in performance for ChemCPA when generalizing to unseen perturbations. As Biolord and scGen cannot generalize to unseen perturbations without modification, our study highlights the need for developing models that can more effectively generalize to unseen combinations of chemical perturbations.

### Defining negative samples for out-of-distribution prediction of protein-peptide binding affinity

Studying T-cell receptors (TCRs) has become crucial in cancer immunotherapy and human infectious disease research [96]. TCRs can detect processed peptides within infected or abnormal cells. Recent studies have focused on predicting TCR-peptide/-pMHC binding using machine or deep learning methods [89, 79]. Many of these studies use data from the Immune Epitope Database (IEDB) [97], VDJdb [98], and McPAS-TCR [99], which predominantly contain CDR3-beta data and lack information on CDR3-alpha. While these methods perform well on test sets from the same source as the training set, they struggle with out-of-distribution samples [91]. This study evaluates cutting-edge methods for out-of-distribution predictions by splitting datasets through a hard split [28]. Additionally, the study reveals a significant sensitivity in model performance to the choice of heuristic for generating negative samples [30], emphasizing the need for further dataset curation and the evaluation of out-of-distribution samples. The datasets have been made available in TDC-2 containing CDR3-beta and CDR3-alpha sequences (Section 7.1).

### Limitations and societal considerations

Open-source datasets and benchmark providers like [17], [35], and [11, 12] play a role in advancing AI by enabling accessible and standardized evaluation methods. However, their limitations and potential negative societal impacts include the risk of biased or incomplete data, which may lead to inaccurate or non-representative AI models. Additionally, the open accessibility of such datasets could lead to misuse, including unethical applications or the proliferation of AI models that reinforce existing biases in medicine. Moreover, reliance on standardized benchmarks may discourage innovation and lead to the over-fitting of models to specific datasets, potentially limiting their generalization in real-world scenarios. Last, evaluating deep learning models for genetic perturbation tasks requires reconsideration in light of recent findings that question their effectiveness for this problem. A recent study revealed that deep learning models do not consistently outperform simpler linear models across various benchmarks [100]. TDC-2’s datasets, benchmarks, metrics, and foundation model tooling lay the foundation for more thorough study, development, and evaluation of models in this space.

## 5 Conclusion

TDC-2 introduces an API-first-dataset architecture that supports retrieving continually updated heterogeneous data sources. The architecture provides abstractions for complex data processing pipelines [11], data lineage [23], and versioning [24]. It augments the stability of emerging biomedical AI workflows, based on advancements from [20, 21, 101], with continuous data updates [25]. It does so via the development of a multi-modal data and model retrieval API leveraging the Model-View-Controller [33, 102, 103] paradigm to introduce data views [104] and a domain-specific-language [105] (section 6.3.1).

The Commons 2.0 (TDC-2) presents a collection of datasets, tools, models, and benchmarks integrating cell-type-specific contextual features with ML tasks across the range of therapeutics. TDC-2 drastically expands the modalities previously available on TDC [11, 12]. TDC-2 supports a far larger set of data modalities and ML tasks than other dataset collections [2] and benchmarks [17, 38, 18, 35]. Modalities in TDC-2 include but are not limited to: single-cell gene expression atlases [2, 51], chemical and genetic perturbations [69], clinical trial data [14], peptide sequence data [90, 91], peptidomimetics protein-peptide interaction data from AS-MS spectroscopy [15, 106], novel 3D structural protein data [107, 108, 109], and cell-type-specific protein-protein interaction networks at single-cell resolution [4]. TDC-2 introduces ML tasks taking on open challenges, including the inferential gap in precision medicine [110, 111] and evaluation on longitudinal data (equation 20), model generalization across cell lines [26, 31] and single-cell perturbations [27] that were not encountered during model training, and evaluation of models across a broad range of diverse biological contexts [4, 30].

TDC-2 is a platform that quantitatively defines open challenges in single-cell therapeutics, determines the current state-of-the-art solutions, promotes method development to improve these solutions, and monitors progress toward these goals. TDC-2 enables broader accessibility for scientists to contribute to advancing the field of single-cell therapeutics. TDC-2 will shift the perspective on method selection and evaluation for therapeutics discovery and machine learning scientists, supporting a transition towards higher standards for methods in contextual AI for therapeutics.

## 6 Appendix

This technical appendix, along with supplementary in section 7, provides a detailed overview of the design, tasks, and benchmarks introduced by TDC-2.

All code and documentation can be found in the TDC-2 Github repository. The URL is https://github.com/mims-harvard/TDC/tree/main. In addition, our website contains all datasets and licenses and further documentation https://tdcommons.ai/.

## 6.1 Data Availability

The website contains all informaton on all datasets discussed in this manuscript under their corresponding tasks. These are: TDC.scDTI (https://tdcommons.ai/multi_pred_tasks/scdti/), TDC.PerturbOutcome (https://tdcommons.ai/multi_pred_tasks/counterfactual/), TDC.TCREpitope (https://tdcommons.ai/multi_pred_tasks/tcrepitope/), TDC.TrialOutcome (https://tdcommons.ai/multi_pred_tasks/trialoutcome/), TDC.SBDD (https://tdcommons.ai/generation_tasks/sbdd/). In addition, all TDC datasets are made available via the harvard dataverse https://dataverse.harvard.edu/dataset.xhtml?persistentId=doi:10.7910/DVN/21LKWG. Instructions for accessing datasets via the TDC Python API can be found in section 7.1.

## 6.2 Code Availability

**All code and documentation can be found in our Github repo**. The URL is https://github.com/mims-harvard/TDC/tree/main.

### 6.3 Methods

### 6.3.1 API-First Design and Model-View-Controller

TDC-2 drastically expands dataset retrieval capabilities available in TDC-1 beyond those of other leading benchmarks. Leading benchmarks, like MoleculeNet [35] and TorchDrug [17] have traditionally provided dataloaders to access file dumps. TDC-2 introduces API-integrated multimodal data-views [33, 112, 104]. To do so, the software architecture of TDC-2 was redesigned using the Model-View-Controller (MVC) design pattern [103, 102]. The MVC architecture separates the model (data logic), view (UI logic), and controller (input logic), which allows for the integration of heterogeneous data sources and ensures consistency in data views [33]. The MVC pattern supports the integration of multiple data modalities by using data mappings and views [104]. The MVC-enabled-multimodal retrieval API is powered by TDC-2’s Resource Model (Section 6.3.2).

#### TDC DataLoader (*Model*)

As per the TDC-1 specification, this component queries the underlying data source to provide raw or processed data to upstream function calls. We augmented this component beyond TDC-1 functionality to allow for querying datasets introduced in TDC-2, such as the CZ CellXGene.

#### TDC meaningful data splits and multimodal data processing (*View*)

As per the TDC-1 specification, this component implements data splits to evaluate model generalizability to out-of-distribution samples and data processing functions for multiple modalities. We augmented this component to act on data views [33] specified by TDC-2’s controller.

#### TDC-2 Domain-Specific Language (*Controller*)

TDC-2 develops an Application-Embedded Domain-Specific Data Definition Programming Language facilitating the integration of multiple modalities by generating data views from a mapping of multiple datasets and functions for transformations, integration, and multimodal enhancements, while mantaining a high level of abstraction [105] for the Resource framework. We include examples developing multimodal datasets leveraging this MVC DSL in listing 2.

### 6.3.2 Resource Model

The Commons introduces a redesign of TDC-1’s dataset layer into a new data model dubbed the TDC-2 resource, which has been developed under the MVC paradigm to integrate multiple modalities into the API-first model of TDC-2.

#### CZ CellXGene with single cell biology datasets

CZ CellXGene [2] is a open-source platform for analysis of single-cell RNA sequencing data. We leverage the CZ CellXGene to develop a TDC-2 Resource Model for constructing large-scale single-cell datasets that maps gene expression profiles of individual cells across tissues, healthy and disease states. TDC-2 leverages the SOMA (Stack of Matrices, Annotated) API, adopts TileDB-SOMA [113] for modeling sets of 2D annotated matrices with measurements of features across observations, and enables memory-efficient querying of single-cell modalities (i.e., scRNA-seq, snRNA-seq), across healthy and diseased samples, with tabular annotations of cells, samples, and patients the samples come from.

We develop a remote procedure call (RPC) API taking the string name (e.g., listing 3 in section 7.3.1) of the desired reference dataset as specified in the CellXGene [2]. The remote procedure call for fetching data is specified as a Python generator expression, allowing the user to iterate over the constructed single-cell atlas without loading it into memory [114]. Specifying the RPC as a Python generator expression allows us to make use of memory-efficient querying as provided by TileDB [113]. The single cell datasets can be integrated with therapeutics ML workflows in TDC-2 by using tools such as PyTorch’s IterableDataset module [115].

#### Knowledge graph, external APIs, and model hub

We have developed a framework for biomedical knowledge graphs to enhance multimodality of dataset retrieval via TDC-2’s Resource Model. Our system leverages PrimeKG to integrate 20 high-quality resources to describe 17,080 diseases with 4,050,249 relationships [22]. Our framework also extends to external APIs, with data views currently leveraging BioPython [116], for obtaining nucleotide sequence information for a given non-coding RNA ID from NCBI [116], and The Uniprot Consortium’s RESTful GET API [117] for obtaining amino acid sequences. In addition we’ve developed the framework to allow access to embedding models under diverse biological contexts via the TDC-2 Model Hub. Examples using these components are in sections 7.3.2 and 7.3.3.

## 6.4 Experiments

### 6.4.1 TDC.TrialOutcome

TDC-2 introduces a model framework, task definition, dataset, and benchmark for the Clinical Outcome Prediction task tailored to precision medicine. The framework and definition aim to assess clinical trials systematically and comprehensively by predicting various endpoints for patient subpopulations. Our benchmark uses the Trial Outcome Prediction (TOP) dataset [14]. TOP consists of 17,538 clinical trials with 13,880 small-molecule drugs and 5,335 diseases. We include the task formulation (section 7.2.4), dataset details 7.1.6, and benchmark (section 7.3.5).

#### Dataset and benchmark

Our benchmark uses the Trial Outcome Prediction (TOP) dataset [14]. TOP consists of 17,538 clinical trials with 13,880 small-molecule drugs and 5,335 diseases. Out of these trials, 9,999 (57.0%) succeeded (i.e., meeting primary endpoints), and 7,539 (43.0%) failed. Out of these trials, 1,787 were in Phase I testing (toxicity and side effects), 6,102 in Phase II (efficacy), and 4,576 in Phase III (effectiveness compared to current standards). We perform a temporal split for benchmarking. The train/validation and test are time-split by the date January 1, 2014, i.e., the start dates of the test set are after January 1, 2014, while the completion dates of the train/validation set are before January 1, 2014. Here, the HINT model [14], is benchmarked against COMPOSE [118] and DeepEnroll [119] models. Results are shown in table 6.

### 6.4.2 Reproducing TDC-2 Benchmarks

Here, we include the instructions for replicating TDC-2 benchmarks, the total amount of computing, and the type of resources used. Code and data details are in sections 7.3.5 and 7.3.4.

**TDC.scDTI**. For benchmarking across ten seeds and another model benchmark, see Section 7.3.5. For pre-training, the best hyperparameters are as follows: the dimension of the nodes’ feature matrix = 1024, dimension of the output layer = 16, lambda = 0.1, learning rate for link prediction task = 0.01, learning rate for protein’s cell type classification task = 0.1, number of attention heads = 8, weight decay rate = 0.00001, dropout rate = 0.6, and normalization layers are layernorm and batchnorm. For pre-training, models are trained on a single NVIDIA Tesla V100-SXM2-16GB GPU. Hyperparameters are used for fine-tuning, as per the Github documentation linked in section 7.3.5. Models are trained on a single NVIDIA Tesla M40 GPU. The relevant function calls are documented in Section 7.3.5.

**Listing 1:**
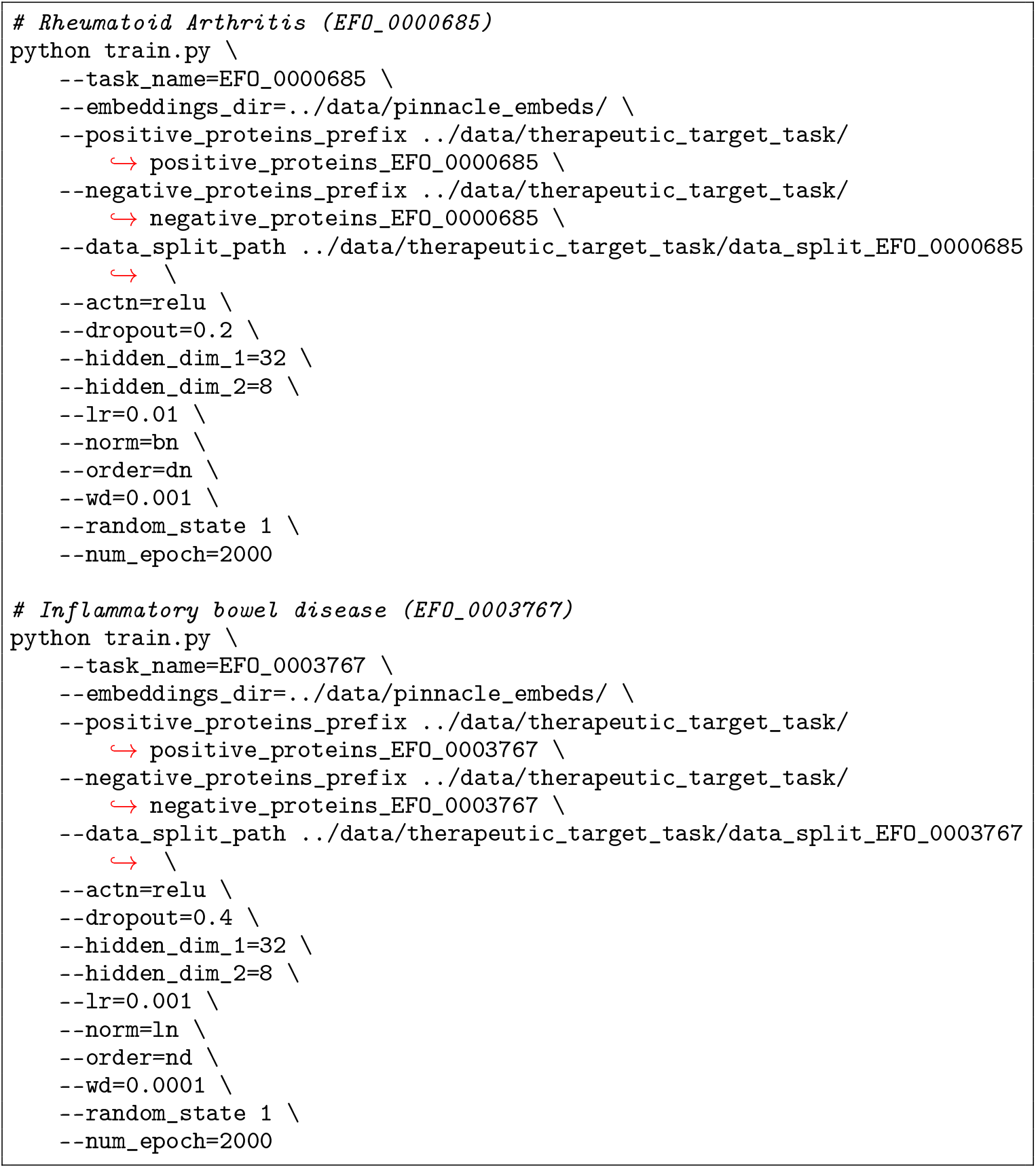
Command line invocations to reproduce the TDC.scDTI benchmark results for [4]. These can also be found by following links in section 7.3.5 or directly in https://github.com/mims-harvard/PINNACLE/tree/main/finetune_pinnacle.

#### TDC.PerturbOutcome – Genetic

All benchmarked methods follow the training procedure described in [26]. Specifically, we use the simulation data split to mimic the real-world use case of genetic perturbation machine learning models. For the Norman double combination perturbation dataset, we withhold perturbations that are either both unseen, one unseen, or both seen in the test set. For the Replogle K562 and RPE1 single perturbation datasets, we split the data by single genes and test on unseen single-gene perturbations. The hyperparameters used were optimal after optimization as reported in [26]. Each model run was executed on an internal high-performance cluster with an Ubuntu 16.04 operating system, using a single Nvidia Quadro RTX 8000 48GB GPU. The code to reproduce the experiment is available at https://github.com/mims-harvard/TDC/tree/main/examples/multi_pred/geneperturb.

#### TDC.PerturbOutcome – Chemical

Benchmark results can be reproduced with code in section 7.3.5. Default settings were used from each model’s GitHub repository, and the experiments were run using Nvidia A100.

#### TDC.TCREpitope

The models for TCR-epitope binding prediction were run on a single A100. We prepared the input data files in the format (most in CSV files) according to the official tutorials. Unknown amino acid letters were replaced by X or removed according to the method requirements. If CDR3A and CDR3B are available, the models will be trained on both unless they can only accept one TCR sequence as input (such as TITAN). If CDR3A is unavailable (ET data), all the models will be trained in the beta-only module. We kept the default parameters to run all the methods. For running TITAN, we transferred the amino acid sequences of epitopes to the SMILE sequences as the inputs. To keep the unseen scenario, we used a zero-shot module of PanPep in the tests of all the data settings. The code for reproducing our benchmark results is in table 7.3.5.

## 7 Supplementary

### 7.1 Datasets

The website tdcommons.ai contains all datasets discussed in this manuscript under their corresponding tasks. These are: TDC.scDTI (https://tdcommons.ai/multi_pred_tasks/scdti/), TDC.PerturbOutcome (https://tdcommons.ai/multi_pred_tasks/counterfactual/), TDC.TCREpitope (https://tdcommons.ai/multi_pred_tasks/tcrepitope/), TDC.TrialOutcome (https://tdcommons.ai/multi_pred_tasks/trialoutcome/), TDC.SBDD (https://tdcommons.ai/generation_tasks/sbdd/). In addition, all TDC datasets are made available via the harvard dataverse https://dataverse.harvard.edu/dataset.xhtml?persistentId=doi:10.7910/DVN/21LKWG. Here we include dataset curation details and code for accessing all datasets for the introduced tasks.

#### 7.1.1 (Li, Michelle, et al.) Dataset

To curate target information for a therapeutic area, we examine the drugs indicated for the therapeutic area of interest and its descendants. The two therapeutic areas examined are rheumatoid arthritis (RA) and inflammatory bowel disease. For rheumatoid arthritis, we collected therapeutic data (i.e., targets of drugs indicated for the therapeutic area) from OpenTargets for rheumatoid arthritis (EFO 0000685), ankylosing spondylitis (EFO 0003898), and psoriatic arthritis (EFO 0003778). For inflammatory bowel disease, we collected therapeutic data for ulcerative colitis (EFO 0000729), collagenous colitis (EFO 1001293), colitis (EFO 0003872), proctitis (EFO 0005628), Crohn’s colitis (EFO 0005622), lymphocytic colitis (EFO 1001294), Crohn’s disease (EFO 0000384), microscopic colitis (EFO 1001295), inflammatory bowel disease (EFO 0003767), appendicitis (EFO 0007149), ulcerative proctosigmoiditis (EFO 1001223), and small bowel Crohn’s disease (EFO 0005629).

We define positive examples (i.e., where the label y = 1) as proteins targeted by drugs that have at least completed phase 2 of clinical trials for treating a specific therapeutic area. As such, a protein is a promising candidate if a compound that targets the protein is safe for humans and effective for treating the disease. We retain positive training examples activated in at least one cell type-specific protein interaction network.

We define negative examples (i.e., where the label y = 0) as druggable proteins that do not have any known association with the therapeutic area of interest according to Open Targets. A protein is deemed druggable if targeted by at least one existing drug. We extract drugs and their nominal targets from Drugbank. We retain negative training examples activated in at least one cell type-specific protein interaction network.

##### Dataset statistics

The final number of positive (negative) samples for RA and IBD were 152 (1,465) and 114 (1,377), respectively. In [4], this dataset was augmented to include 156 cell types.

**Dataset split. Cold Split We split the dataset such that about 80% of the proteins are in the training set, about 10% of the proteins are in the validation set, and about 10% of the proteins are in the test set. The data splits are consistent for each cell type context to avoid data leakage.**

**References**

**Dataset license**. CC BY 4.0

##### Code Sample

The dataset and splits are currently available on TDC Harvard Dataverse. In addition, you may obtain the protein splits used in [4] via the following code.

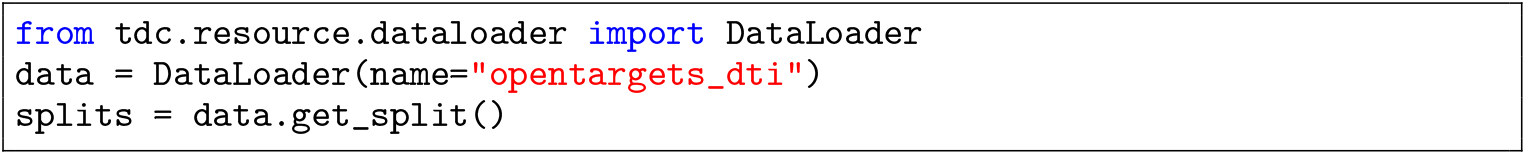

### 7.12 scPerturb Dataset

The scPerturb dataset is a comprehensive collection of single-cell perturbation data harmonized to facilitate the development and benchmarking of computational methods in systems biology. It includes various types of molecular readouts, such as transcriptomics, proteomics, and epigenomics. scPerturb is a harmonized dataset that compiles single-cell perturbation-response data. This dataset is designed to support the development and validation of computational tools by providing a consistent and comprehensive resource. The data includes responses to various genetic and chemical perturbations, crucial for understanding cellular mechanisms and developing therapeutic strategies. Data from different sources are uniformly pre-processed to ensure consistency. Rigorous quality control measures are applied to maintain high data quality. Features across different datasets are standardized for easy comparison and integration.

#### Dataset statistics

44 publicly available single-cell perturbation-response datasets. Most datasets have, on average, approximately 3000 genes measured per cell. 100,000+ perturbations.

**Dataset split. Cold Split and Random Split defined on cell lines and perturbation types.**

**References[69]**

**Dataset license. CC BY 4.0**

#### Code Sample

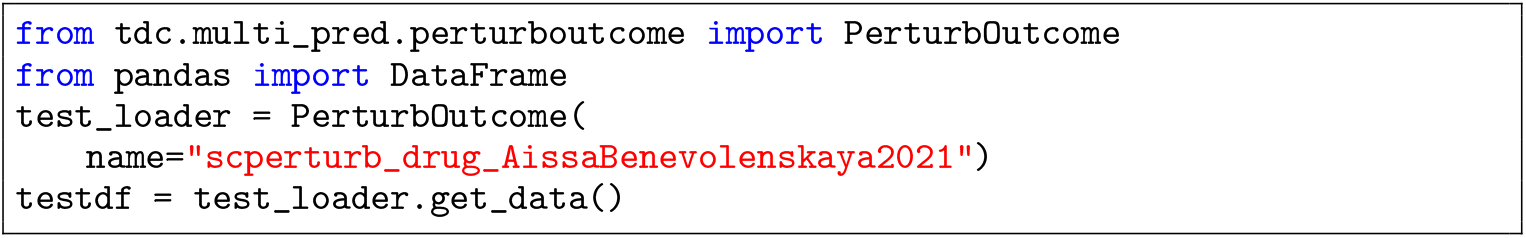

#### 7.1.3 TCHard Dataset

The TChard dataset is designed for TCR-peptide/-pMHC binding prediction. It includes over 500,000 samples from sources such as IEDB, VDJdb, McPAS-TCR, and the NetTCR-2.0 repository. The dataset is utilized to investigate how state-of-the-art deep learning models generalize to unseen peptides, ensuring that test samples include peptides not found in the training set. This approach highlights the challenges deep learning methods face in robustly predicting TCR recognition of peptides not previously encountered in training data.

##### Dataset statistics. 500,000 samples

**Dataset split**. Cold Split referred to as “Hard” split in [90].

**References [90]**

**Dataset license**. Non-Commercial Use

##### Code Sample

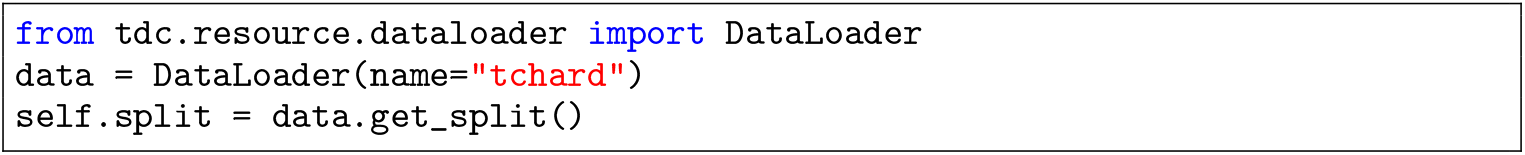

#### 7.1.4 PanPep Dataset

PanPep is a framework constructed in three levels for predicting the peptide and TCR binding recognition. We have provided the trained meta learner and external memory, and users can choose different settings based on their data available scenarios: Few known TCRs for a peptide: few-shot setting; No known TCRs for a peptide: zero-shot setting; plenty of known TCRs for a peptide: majority setting. More information is available in the Github repo.

##### Dataset statistics. Data from multiple studies involving millions of TCR sequences

**Dataset split. Cold Split** referred to as “Hard” split in [91].

**References [91]**

**Dataset license**. GPL-3.0

##### Code Sample

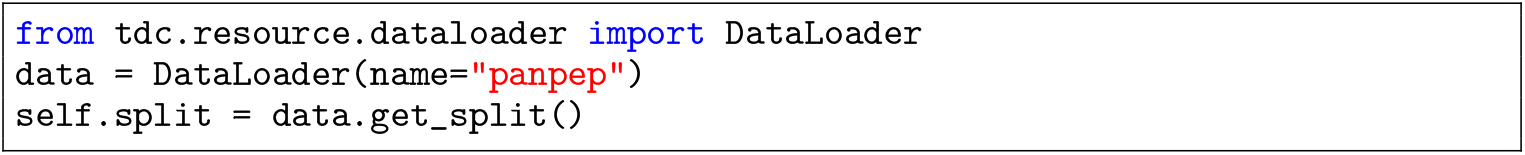

#### 7.1.5 (Ye X et al) Dataset

Affinity selection-mass spectrometry data of discovered ligands against single biomolecular targets (MDM2, ACE2, 12ca5) from the Pentelute Lab of MIT This dataset contains affinity selection-mass spectrometry data of discovered ligands against single biomolecular targets. Several AS-MS-discovered ligands were taken forward for experimental validation to determine the binding affinity (KD) as measured by biolayer interferometry (BLI) to the listed target protein. If listed as a “putative binder,” AS-MS alone was used to isolate the ligands to the target, with KD < 1 uM required and often observed in orthogonal assays, though there is some (< 50%) chance that the ligand is nonspecific. Most of the ligands are putative binders, with 4446 total provided. For those characterized by BLI (only 34 total), the average KD is 266 ± 44 nM; the median KD is 9.4 nM.

##### Dataset statistics. 34 positive ligands, 4446 putative binders, and three proteins

**Dataset Split. Stratified Split** and **N/A Split**: We provide stratified 10/90 split on train/test as well as “test set only” split.

**References [106, 15]**

**Dataset license**. CC BY 4.0

##### Code Sample

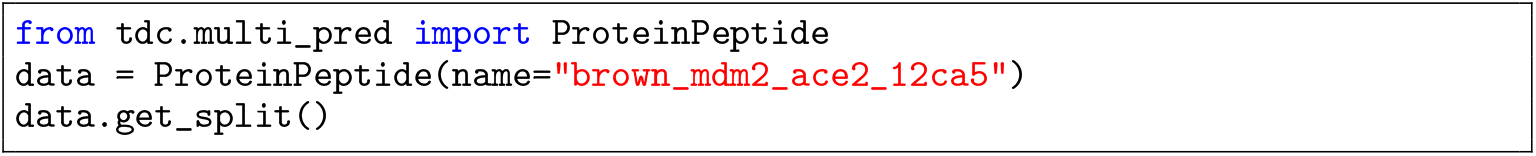

#### 7.1.6 TOP Dataset

TOP [14] consists of 17,538 clinical trials with 13,880 small-molecule drugs and 5,335 diseases. Out of these trials, 9,999 (57.0%) succeeded (i.e., meeting primary endpoints), and 7,539 (43.0%) failed. For each clinical trial, we produce the following four data items: (1) drug molecule information, including Simplified Molecular Input Line Entry System (SMILES) strings and molecular graphs for the drug candidates used in the trials; (2) disease information including ICD-10 codes (disease code), disease description, and disease hierarchy in terms of CCS codes (https://www.hcup-us.ahrq.gov/toolssoftware/ccs10/ccs10.jsp); (3) trial eligibility criteria are in unstructured natural language and contain inclusion and exclusion criteria; and (4) trial outcome information includes a binary indicator of trial success (1) or failure (0), trial phase, start and end date, sponsor, and trial size (i.e., number of participants).

**Dataset statistics**. Phase I: 2,402 trials / Phase II: 7,790 trials / Phase III: 5,741 trials.

**Dataset split. Temporal Split** as defined in [14] and Section 6.4.1.

**References [14]**

**Dataset license**. Non-Commercial Use

##### Code Sample

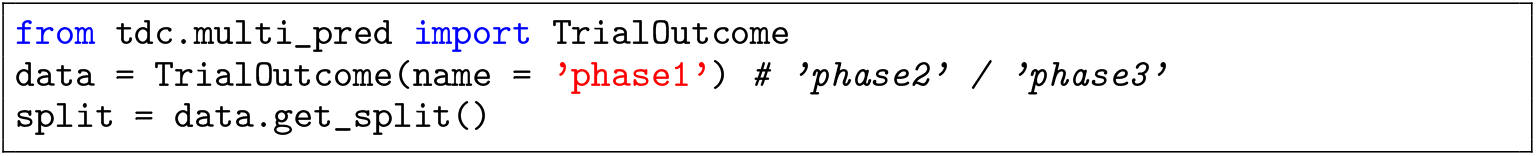

#### 7.1.7 PDBBind Dataset

PDBBind is a comprehensive database extracted from PDB with experimentally measured binding affinity data for protein-ligand complexes. PDBBind does not allow the dataset to be re-distributed in any format. Thus, we could not host it on the TDC server. However, we provide an alternative route since significant processing is required to prepare the dataset ML. The user only needs to register at http://www.pdbbind.org.cn/, download the raw dataset, and then provide the local path. TDC will then automatically detect the path and transform it into an ML-ready format for the TDC data loader.

**Dataset statistics**. 19,445 protein-ligand pairs

**Dataset split. Random Split**

**References [107]**

**Dataset license**. See note in the description on the TDC website.

*Code Sample*

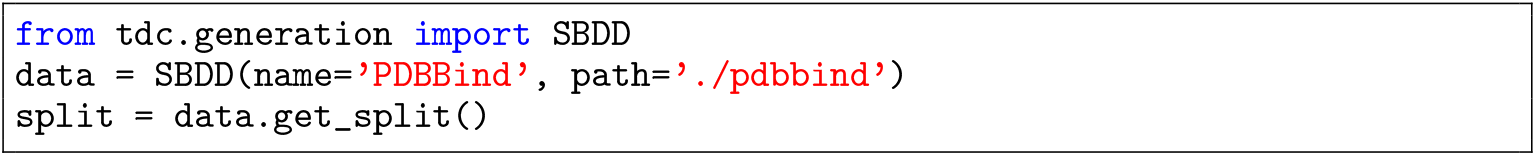

**Dataset statistics**. 22,886 active compounds and affinities against 102 targets. DUD-E does not support pocket extraction as protein and ligand are not aligned.

**Dataset split. Random Split**

**References [109]**

**Dataset license**. Not specified

##### Code Sample

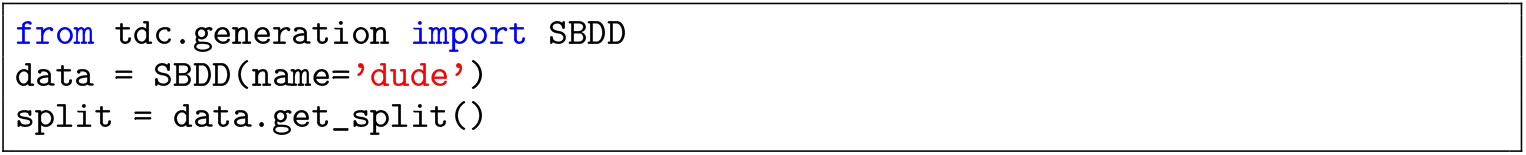

#### 7.1.8 scPDB Dataset

scPDB is processed from PDB for structure-based drug design that identifies suitable binding sites for protein-ligand docking.

##### Dataset statistics. 16,034 protein-ligand pairs over 4,782 proteins and 6,326 ligands

**Dataset split. Random Split**

**References [108]**

**Dataset license**. Not specified

##### Code Sample

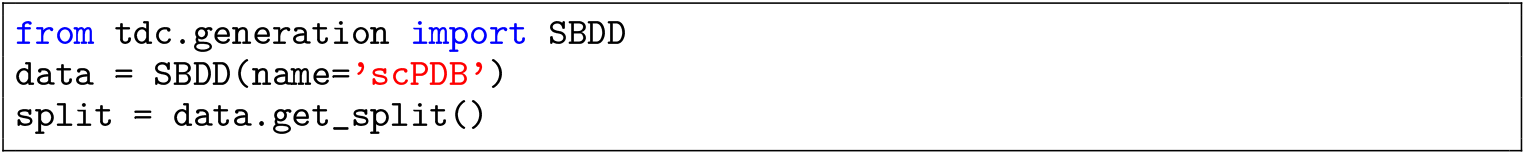

### 7.2 Equations

#### 7.2.1 TDC.scDTI: Contextualized Drug-Target Nomination (Identification)

TDC-2 introduces TDC.scDTI task. The predictive, non-generative task is formalized as learning an estimator for a disease-specific function *f* of a target protein and cell type outputting whether the candidate protein t is a therapeutic target in that cell type *c*:

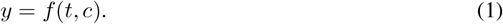

##### Target candidate set

The target candidate set includes proteins, nucleic acids, or other molecules drugs can interact with, producing a therapeutic effect or causing a biological response. The target candidate set is constrained to proteins relevant to the disease being treated. It is denoted by:

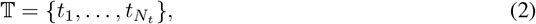

where 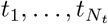 are *N*_*t*_ target candidates for the drugs treating the disease. Information modeled for target candidates can include interaction, structural, and sequence information.

##### Biological context set

The biological context set includes the cell-type-specific contexts in which the target candidate set operates. This set is denoted as:

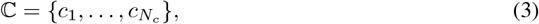

where 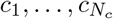 are *N*_*c*_ biological contexts on which drug-target interactions are being evaluated. Information modeled for cell-type-specific biological contexts can include gene expression and tissue hierarchy. The set is constrained to disease-specific cell types and tissues.

**Drug-target identification**. Drug-Target Identification is a binary label *y ∈ {* 1, 0 *}*, where *y* = 1 indicates the protein is a candidate therapeutic target. At the same time, 0 means the protein is not such a target.

The goal is to train a model *f*_*θ*_ for predicting the probability ŷ*∈* [0, 1] that a protein is a candidate therapeutic target in a specific cell type. The model learns an estimator for a disease-specific function of a protein target *t ∈* 𝕋 and a cell-type-specific biological context *c∈* ℂ as input, and the model is tasked to predict:

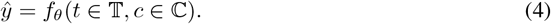

#### 7.2.2 TDC.PerturbOutcome: Perturbation-Response Problem Formulation

TDC-2 introduces Perturbation-Response prediction task. The predictive, non-generative task is formalized as learning an estimator for a function of the cell-type-specific gene expression response to a chemical or genetic perturbation, taking a perturbation *p ∈* ℙ, a pre-perturbation gene expression profile from the control set e_0_ *∈* 𝔼_⊬_, and the biological context *c∈* ℂ under which the gene expression response to the perturbation is being measured:

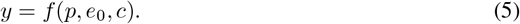

We center our definition on regression for the cell-type-specific gene expression vector in response to a chemical or genetic perturbation.

**Perturbation set**. The perturbation set includes genetic and chemical perturbations. It is denoted by:

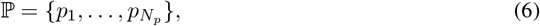

where 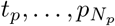 are *N*_*p*_ evaluated perturbations. Information modeled for genetic perturbations can include the type of perturbation (i.e., knockout, knockdown, overexpression) and target gene(s) of the perturbation. Information modeled for chemical perturbations can include chemical structure (i.e., SMILES, InChl) and concentration and duration of treatment.

**Control set**. The control set includes the unperturbed gene expression profiles. This set is denoted as:

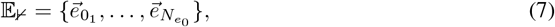

where 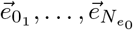 are 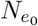 unperturbed gene expression profile vectors. Information models for gene expression profiles can include raw or normalized gene expression counts, transcriptomic profiles, and isoform-specific expression levels.

**Biological context set**. The biological context set includes the cell-type-specific contexts under which the perturbed gene expression profile is measured. It is denoted by:

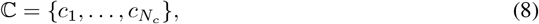

where 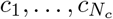 are the *N*_*c*_ biological contexts under which perturbations are being evaluated. Information modeled for biological contexts can include cell type or tissue type and experimental conditions [69] as well as epigenetic markers [121, 122].

##### Perturbation-response readouts

Perturbation-Response is a gene expression vector 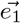, where 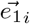 denotes the expression of the i-th gene in the vector. It is the outcome of applying a perturbation, p_*i*_ *∈* ℙ, within a biological context, c_*j*_ *∈* ℂ, to a cell with a measured control gene expression vector, 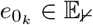.

The Perturbation-Response Prediction learning task is to learn a regression model *f*_*θ*_ estimating the perturbation-response gene expression vector 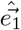 for a perturbation applied in a cell-type-specific biological context to a control:

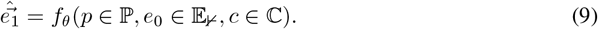

#### 7.2.3 TDC.ProteinPeptide: Protein-Peptide Interaction Prediction Problem Formulation

TDC-2 introduces the Protein-Peptide Binding Affinity prediction task. The predictive, non-generative task is to learn a model estimating a function of a protein, peptide, antigen processing pathway, biological context, and interaction features. It outputs a binding affinity value (e.g., dissociation constant Kd, Gibbs free energy ΔG) or binary label indicating strong or weak binding. The binary label can also include additional biomarkers, such as allowing for a positive label if and only if the binding interaction is specific [15, 123, 124]. To account for additional biomarkers beyond binding affinity value, our task is specified with a binary label.

**Protein set**. The protein set includes target proteins. It is denoted by:

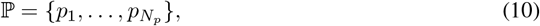

where 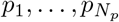 are *N*_*p*_ target proteins. Information modeled for proteins can include sequence, structural, or post-translational modification data.

**Peptide set**. The control set includes the peptide candidates. This set is denoted as:

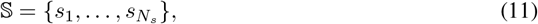

where 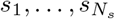 are N_*s*_ candidate peptides. Information modeled for candidate peptides can include sequence, structural, and physicochemical data.

#### Antigen processing pathway set

The antigen processing pathway set includes antigen processing pathway profile information about prior steps in the biological antigen presentation pathway processes. It is denoted by:

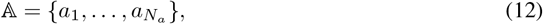

where 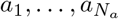 are the *N*_*a*_ antigen processing pathway profiles modeled. Information modeled in a profile can include proteasomal cleavage sites [125], classification into viral, bacterial, and self-protein sources and endogenous vs exogenous processing pathway [126, 127, 84, 128], and target/receptor-specific pathway attributes such as transporter associated with antigen processing (TAP) affinity [129], and endosomal/lysosomal processing efficiency [130].

**Interaction set**. It contains the interaction feature profiles. The set is denoted by:

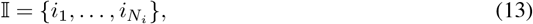

where 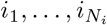 are the *N*_*i*_ interaction feature profiles. Information modeled in an interaction feature profile can include contact maps [131, 132, 133, 134], distance maps [132, 135], electrostatic interactions [131], and hydrogen bonds [131].

#### Cell-type-specific biological context set

It contains the interaction feature profiles. The set is denoted by:

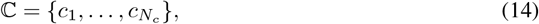

where 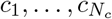 are the *N*_*c*_ cell-type-specific biological contexts under which the protein-peptide interaction is being evaluated. Information modeled in the cell-type-specific biological context can include transcriptomic and proteomic data. We note, however, that, to our knowledge, single-cell transcriptomic and proteomic data has yet to be used in protein-peptide binding affinity prediction, outlining a promising avenue of research in developing machine learning models for peptide-based therapeutics.

**Protein-peptide interaction**. It is a binary label, *y ∈ {*1, 0 *}*, where *y* = 1 indicates a protein-peptide pair met the target biomarkers and *y* = 0 indicates the pair did not meet the target biomarkers.

The Protein-Peptide Interaction Prediction learning task is to learn a binary classification model *f*_*θ*_ estimating the probability, ŷ, of a protein-peptide interaction meeting specific biomarkers:

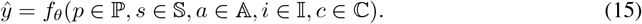

#### 7.2.4 Clinical Trial Outcome Prediction Problem Formulation

The Clinical Trial Outcome Prediction task is formulated as a binary classification problem, where the machine learning model predicts whether a clinical trial will have a positive or negative outcome. It is a function that takes patient data, trial design, treatment characteristics, disease, and macro variables as inputs and outputs a trial outcome prediction, a binary indicator of trial success (1) or failure (0).

**Patient set**. The patient set includes one or multiple patient sub-populations, with the extreme case representing personalization. It is denoted as follows:

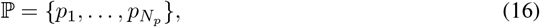

where 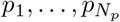 are *N*_*p*_ patient sub-populations in this trial. The TOP benchmark [14] dataset represents patient data as part of the trial eligibility criteria. Patient data can include demographics [136, 137, 138, 139, 140], baseline health metrics [139, 140, 141], and medical history [136, 137, 138, 139, 140].

**Trial design set**. The trial design set includes this clinical trial’s design profiles. It is denoted as:

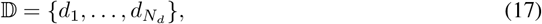

where 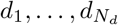 are *N*_*d*_ eligible trial design profiles for this clinical trial. Trial design profiles can model information including phase of the trial [14], number of participants, duration of the trial, trial eligibility criteria [14], and randomization and blinding methods [142, 143, 144].

**Treatment set**. The treatment set includes the candidate treatments for the trial. It is denoted as:

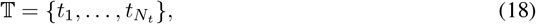

where 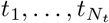 are *N*_*t*_ candidate treatments for the clinical trial. The information modeled for treatments can include type of treatment (drug [14, 145], device [146, 147, 148], procedure [149, 150, 151, 152, 153]), dosage and administration route [142, 141, 154], mechanism of action [155, 156, 157], pre-clinical and early-phase trial results [156, 141, 158, 159].

#### Macro context set

The macro context set contains the configurations of macro variables relevant to the clinical trial. It is denoted as:

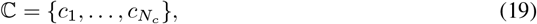

where 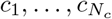 are *N*_*c*_ configurations containing the values for macro variables relevant to the trial, which can include geography [160, 156, 159, 161] and regulatory considerations [156, 160].

**Trial outcomes**. The trial outcome is a binary label *y ∈ {* 1, 0 *}*, where y = 1 indicates the trial met their primary endpoints, while 0 means failing to meet with the primary endpoints.

The learning task is to learn a model *f*_*θ*_ for predicting the trial success probability *ŷ*, where ŷ *∈* [0, 1]:

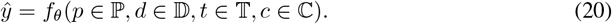

#### 7.2.5 Structure-Based Drug Design Problem Formulation

Structure-based Drug Design aims to generate diverse, novel molecules with high binding affinity to protein pockets (3D structures) and desirable chemical properties. These properties are measured by oracle functions. A machine learning task first learns the molecular characteristics given specific protein pockets from a large set of protein-ligand pair data. Then, from the learned conditional distribution, we can sample novel candidates.

##### Target candidate set

The target candidate set includes proteins, nucleic acids, or other biomolecules drugs can interact with, producing a therapeutic effect or causing a biological response. It is denoted by:

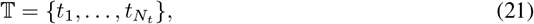

where 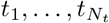 are *N*_*t*_ target candidates for the evaluated set of drugs. Information modeled for target candidates can include interaction, structural, and sequence information.

##### Ligand candidate set

The ligand drug candidate set includes the drug molecules being tested for a particular therapeutic effect or biological response. It is denoted by:

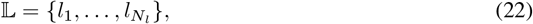

where 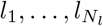 are the N_*l*_ ligand/drug molecules being evaluated. Drug modeling can include molecular structure, often represented in formats such as SMILES (Simplified Molecular Input Line Entry System) or InChI (International Chemical Identifier) [162], physicochemical properties like hydrophobicity and molecular weight [80], and molecular descriptors and fingerprints [163].

**Scoring function**. The scoring function, denoted by S, evaluates the binding affinity of ligand l *∈* 𝕃?to protein target t *t ∈ 𝕋*.

**Drug-likeness function**. Function representing the drug-likeness of ligand *l∈* 𝕃, including properties like solubility, stability, and toxicity.

The generative learning task is to generate the ligand *l ∈* 𝕃 maximizing binding affinity, S_*θ*_, and drug-likeness, *f*_*θ*_. Given a loss function, Loss(*S*(*t, l*), *f*(*l*)), for *t ∈* 𝕋 and *l∈* 𝕃, the first step is to learn a model M_*θ*_ s.t.,

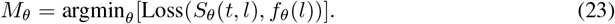

This is followed by the ligand optimization step, which optimizes the ligand for maximum binding affinity and drug-likeness given the trained model. A ligand optimization function, *F*, such as addition or multiplication, is used for the optimization:

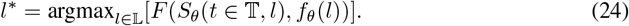

An example formulation would be as follows:

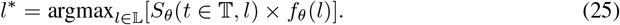

#### 7.2.6 Context-Specific Metrics

Context-specific metrics are defined to measure model performance at critical biological slices, with our benchmarks focused on measuring cell-type-specific model performance. For single-cell drug-target nomination, we measure model performance at top-performing cell types. The metrics chosen were: APR@5 Top-20 CT - average precision and recall at k = 5 for the 20 best-performing cell types (CT); AUROC Top-1 CT - AUROC for top-performing cell type; AUROC Top-10 CT and AUROC Top-20 CT -weighted average AUROC for top-10 and top-20 performing cell types, respectively, each weighted by the number of samples in each cell type. Formally, we define context-specific APR@5 and AUROC below.

##### Context-specific AUROC

To calculate the **AUROC for the top K performing cell types**, we first need to determine which cell types achieve the highest AUROC scores. After selecting the top-performing cell types, we weigh each top-performing cell type’s AUROC score by the number of samples in that cell type.

We denote:

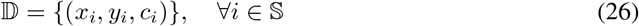

Here, 𝔻 denotes the dataset where x_*i*_ denotes the feature vector, *y*_*i*_ is the true label, and c_*i*_ is the cell type for sample i from 𝕊. We further denote *C*, the set of unique cell types. Then, the AUROC for a specific cell type, *AUROC*_*c*_, is computed as:

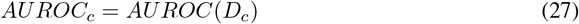

Here, D_*c*_ = *{*(*x*_*i*_, *y*_*i*_) | *c*_*i*_*}* = c is the subset of the dataset for cell type c and AUROC(D_*c*_) represents the AUROC score computed over this subset. Once these are computed, values can be sorted in descending order to select the top X cell type with highest AUROC value.

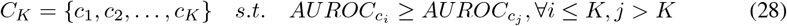

The weighted AUROC for the top K cell types is given by weighting each cell type’s AUROC by the proportion of its samples relative to the total samples in the top K cell types.

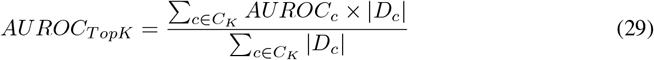

This measure represents a balance between representation and performance of the cell types.

##### Context-specific Average Precision at rank R (AP@R)

In our study, we let R = 5 and compute **AP@5 for the top K performing cell types**. We denote dataset and samples as above.

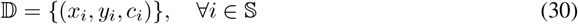

Here, 𝔻 denotes the dataset where x_*i*_ denotes the feature vector, *y*_*i*_ is the true label, and c_*i*_ is the cell type for sample i from 𝕊. We further denote *C*, the set of unique cell types. The samples of each cell type, *D*_*c*_ = (*x*_*i*_, y_*i*_) | *c*_*i*_ = *c*, can be sorted based on the score output by the model for said sample *f*(*x*_*i*_), with average precision at rank type computed accordingly.

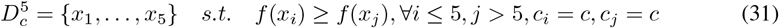

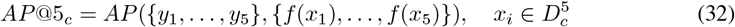

The average precision at rank k at Top X cell types can then be defined as:

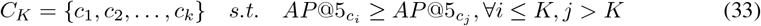

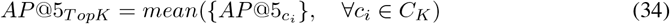

AP summarizes a precision-recall curve as the weighted mean of precisions achieved at each threshold, with the increase in recall from the previous threshold used as the weight. Some key advantages of using AP@K include robustness to (1) varied numbers of protein targets activated across cell type-specific protein interaction networks and (2) varied sizes of cell type-specific protein interaction networks [4]. We compute AP using the scikit package as specified in https://scikit-learn.org/1.5/modules/generated/sklearn.metrics.average_precision_score.html.

### 7.3 Algorithms, Program codes and Listings

We provide code samples for the components described in sections 6.3.1 and 6.3.2.

**Listing 2:**
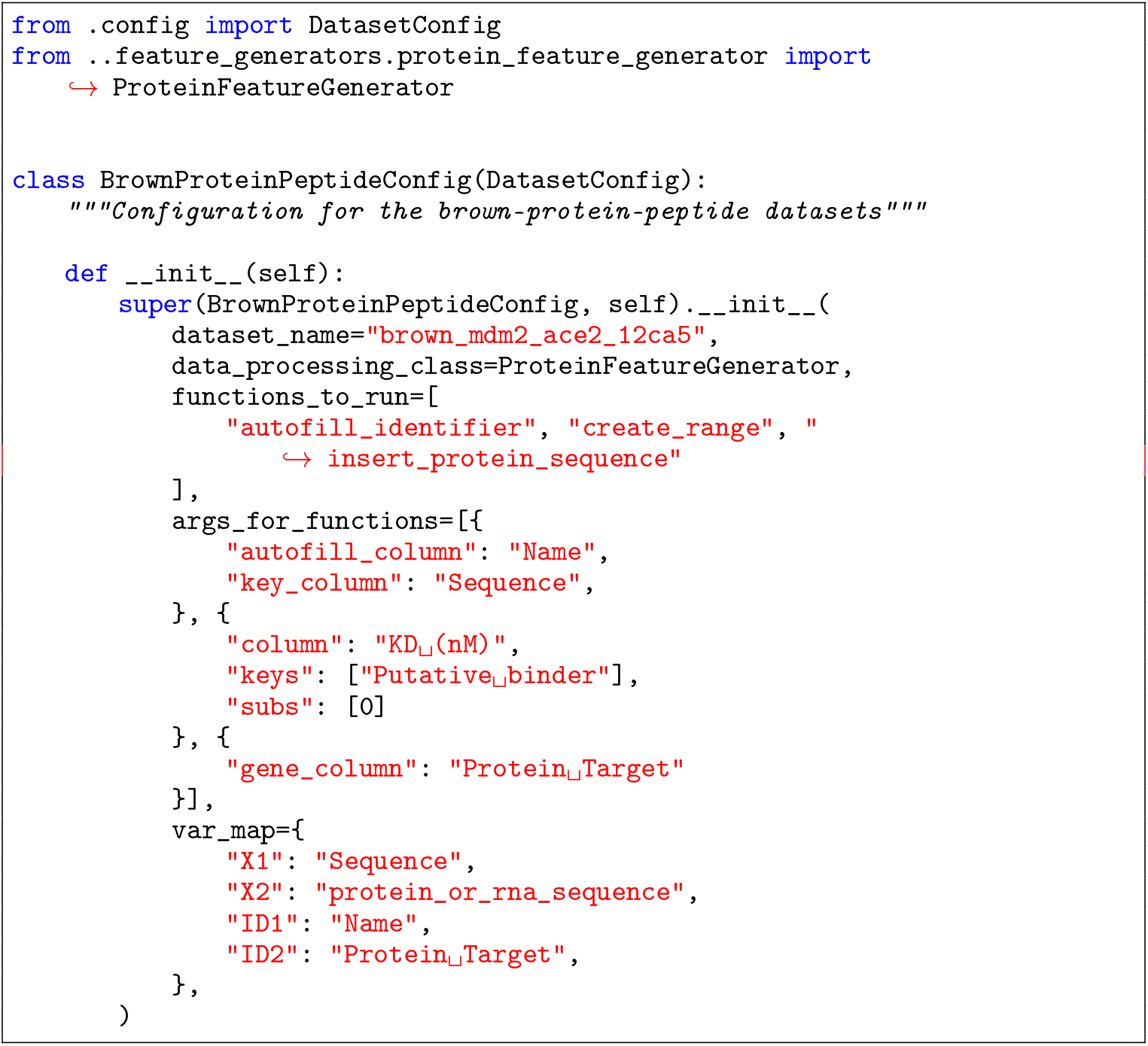
The above configuration augments a protein-peptide dataset with an additional modality, amino acid sequence, and invokes numerous data processing functions tailored to the specific needs of the underlying dataset. Added information for this demonstration can be found at: https://colab.research.google.com/drive/13MYlg5tWpywWbKYsJQXafKAlVF2hz-sP?usp=sharing. There are more complex workflows implemented for current TDC dataviews and all such views leveraging the DSL can be found in the repo at https://github.com/mims-harvard/TDC/blob/main/tdc/dataset_configs/config_map.py

#### 7.3.1 TDC-2 Multimodal Single-Cell Retrieval API

We focus on the use case of an ML researcher who wishes to train a model on a large-scale single-cell atlas. In particular, researchers would be familiar with and have trained models on traditional single-cell datasets such as Tabula Sapiens [51]. Their interest is to scale a model by training it on a more extensive single-cell atlas based on this reference dataset. We build such an API. Specifically, given a reference dataset available in CellXGene Discover [2], we allow the user to perform a memory-efficient query using TileDB-SOMA to expand the reference dataset to include cell entries with non-zero readouts for any of the genes present in the reference dataset. This allows users to build large-scale single-cell atlases on familiar reference datasets. The example below illustrates how a user may construct a large-scale atlas with Tabula Sapiens as the reference dataset. Other use cases include augmenting datasets using knowledge graphs and cell-type-specific biomedical contexts. These capabilities are all powered by the Model-View-Controller framework (section 6.3.1).

**Listing 3:**
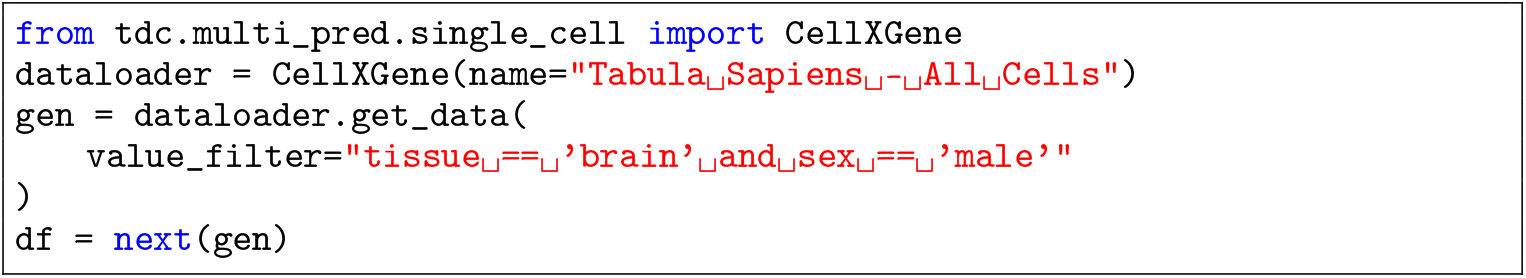
The example below illustrates how a user may construct a large-scale atlas with Tabula Sapiens as the reference dataset using the TDC-2 CELLXGENE API.

**Listing 4:**
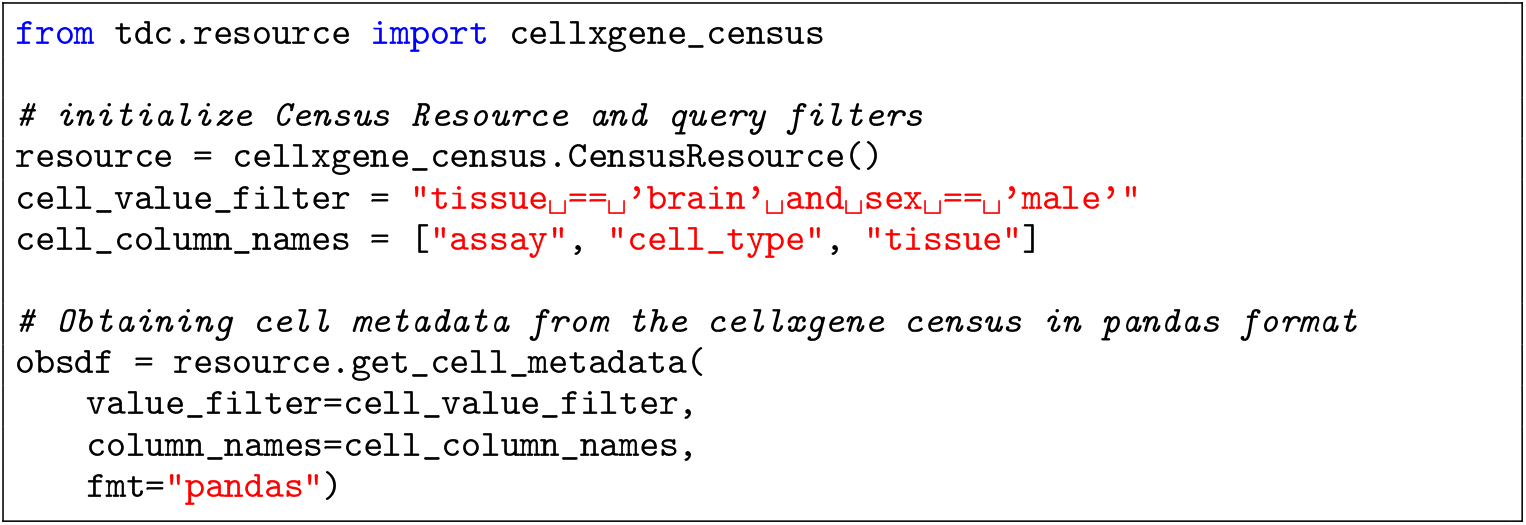
In addition to our TDC-2 DataLoader API implementation for the CellXGene RPC API, we provide a simplified wrapper over the CellXGene Census Discovery API, which allows users to perform remote procedure calls to fetch Cell Census data in more machine-learning-friendly formats like Pandas and Scipy. We also maintain support for the AnnData format. Users can query Cell Census counts as well as metadata using this API. The code sample below illustrates such usage.

#### 7.3.2 PrimeKG Knowledge Graph

PrimeKG supports drug-disease prediction by including an abundance of ‘indications,’ ‘contradictions’, and ‘off-label use’ edges, which are usually missing in other knowledge graphs. We accompany PrimeKG’s graph structure with text descriptions of clinical guidelines for drugs and diseases to enable multimodal analyses [22]. The code below depict example use cases of the TDC-2 PrimeKG API. Demonstrations are additionally available in https://colab.research.google.com/drive/1kYH8nt3nW7tXYBPNcfYuDbWxGTqOEnWg?usp=sharing.

**Listing 5:**
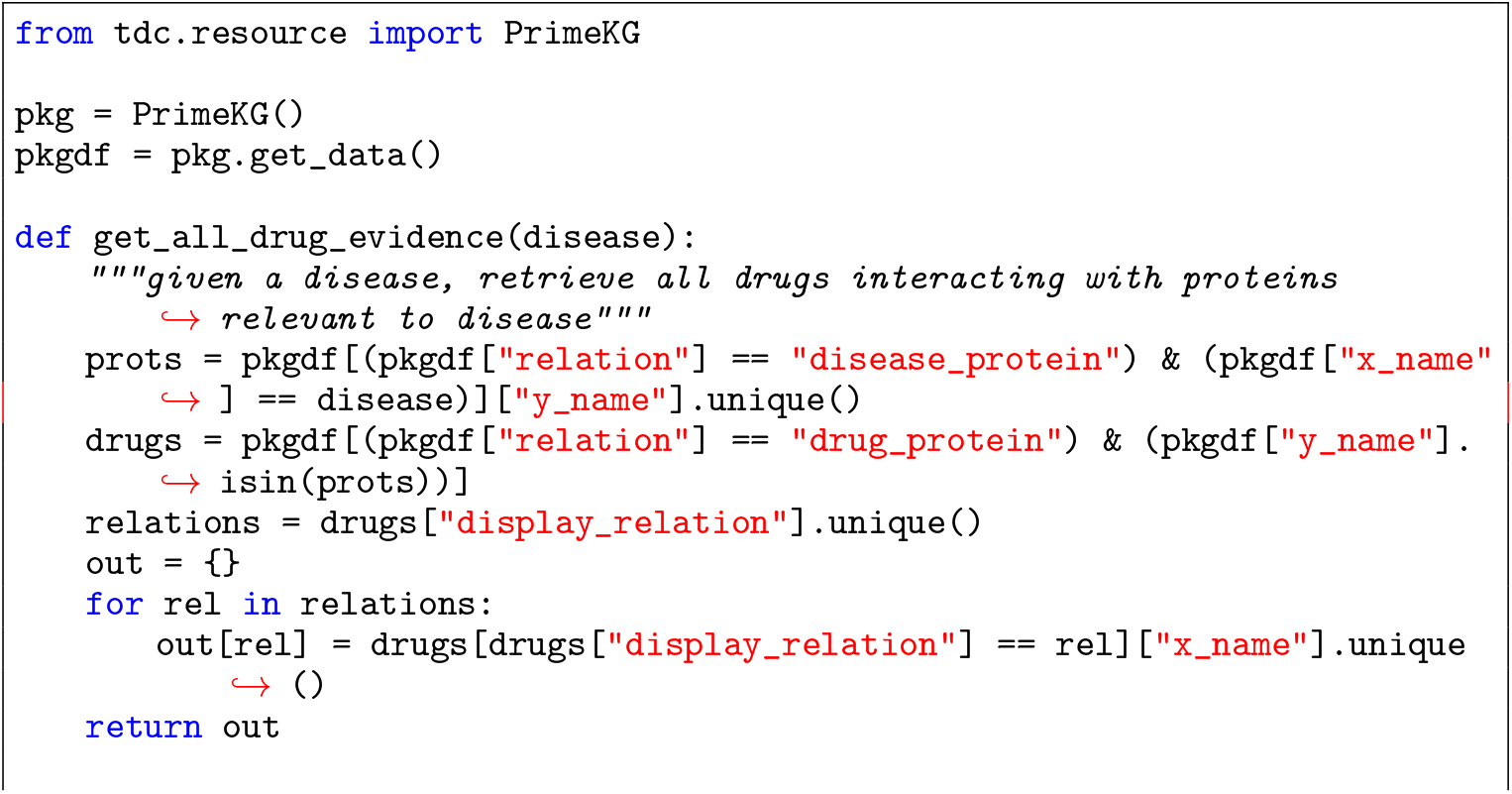
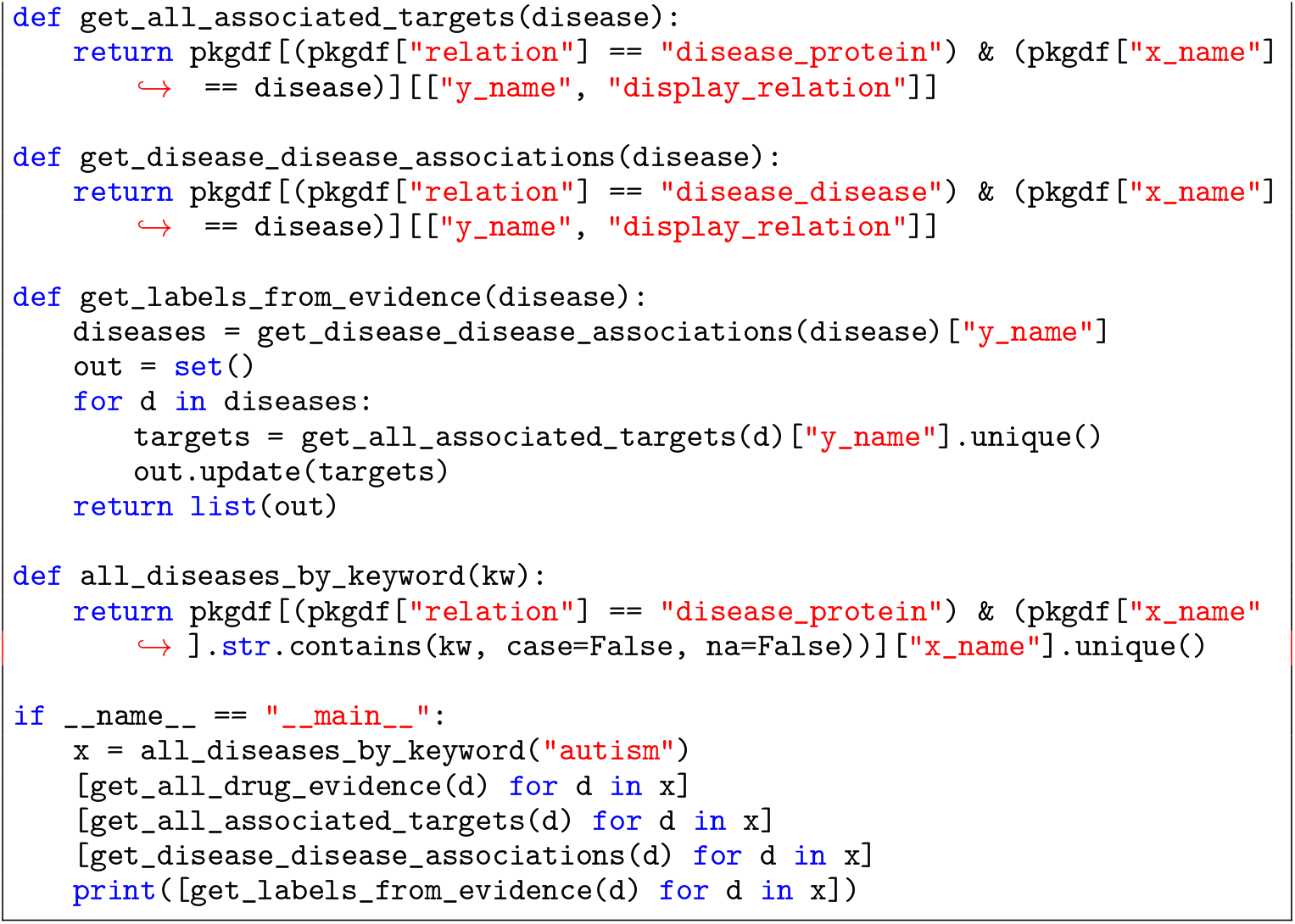
We illustrate here example utilities for retrieving drug-target-disease associations using the TDC-2 PrimeKG API

**Listing 6:**
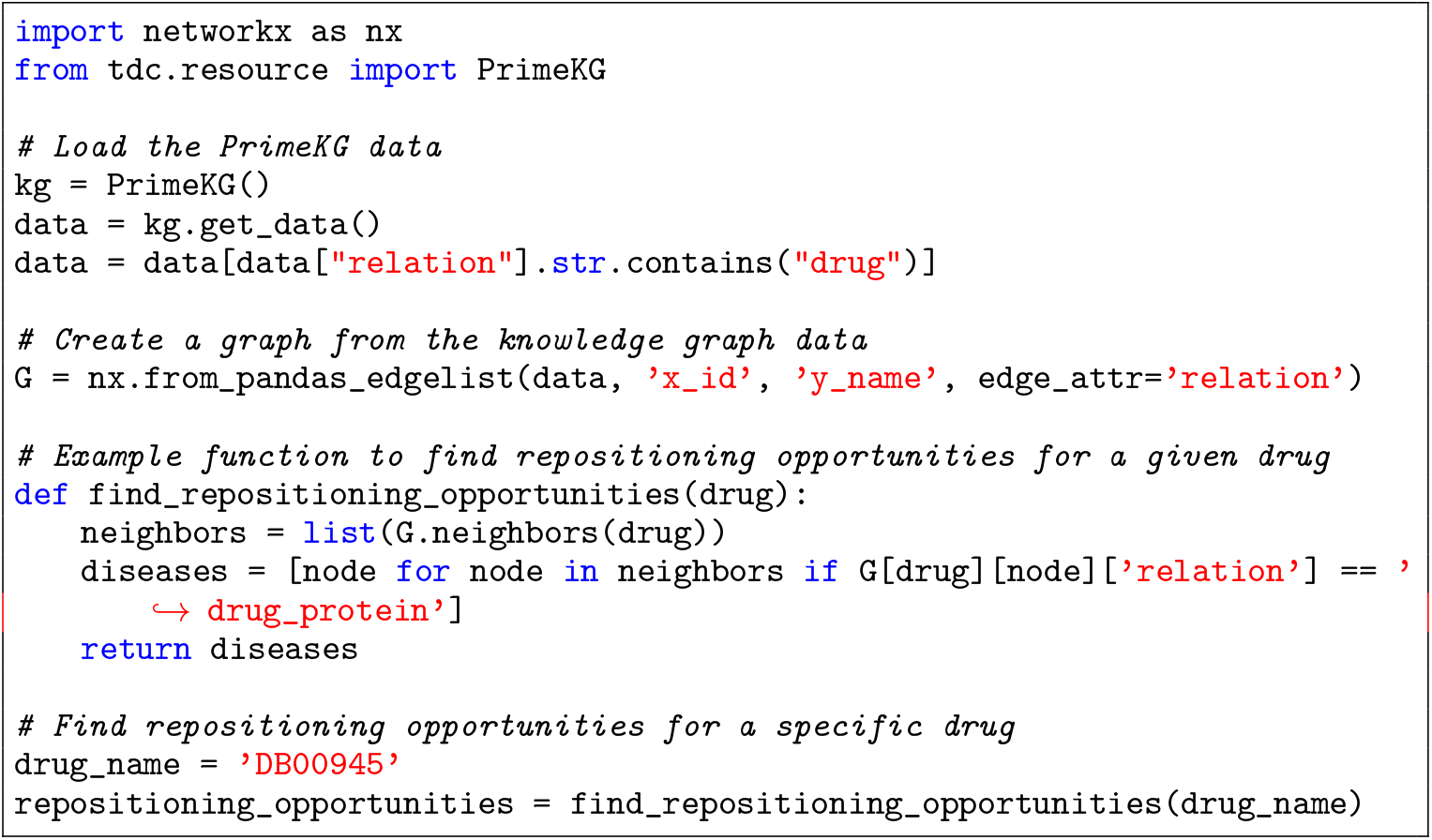
Here we illustrate combinig the TDC-2 PrimeKG API with the networkx module to retrieve drug repositioning opportunities.

#### 7.3.3 TDC-2 Model Server

The introduced model server is composed of the TDC-2 Model Hub and a set of utilities and endpoints for facilitating model inference and fine-tuning. TDC-2 introduces The Commons’ HuggingFace Model Hub. It is a resource with pre-trained models, including geometric deep learning models, large language models, and other contextualized multimodal models for therapeutic tasks. The models can be fine-tuned using datasets in TDC-2 and be used for downstream tasks such as implementations of multi-agent collaborative schemes [164] (i.e., expert consultants) our predictive therapeutic tasks [3, 19]. The model hub details and available models can be found at https://huggingface.co/tdc.

**Listing 7:**
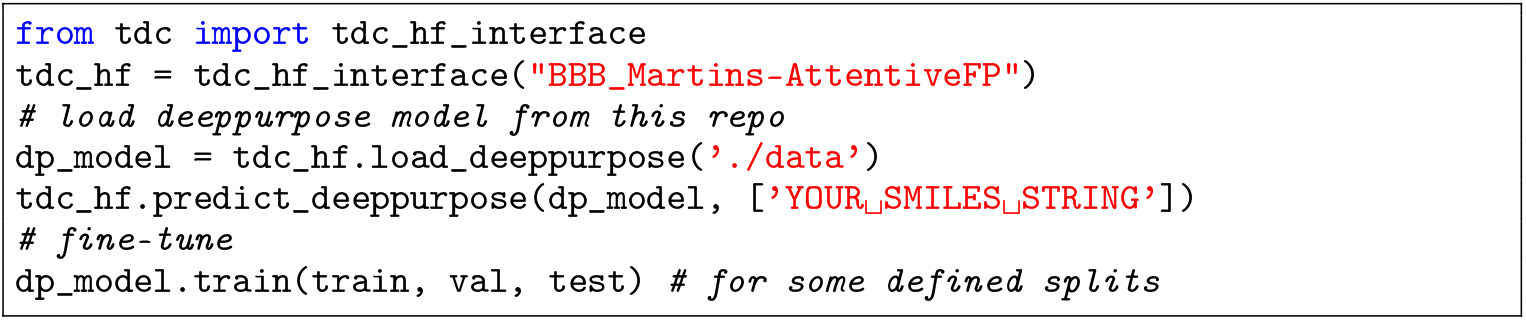
The below illustrates the basic functionality of the model hub to download a model and perform inference on a precdictive task as well as fine-tune the model

**Listing 8:**
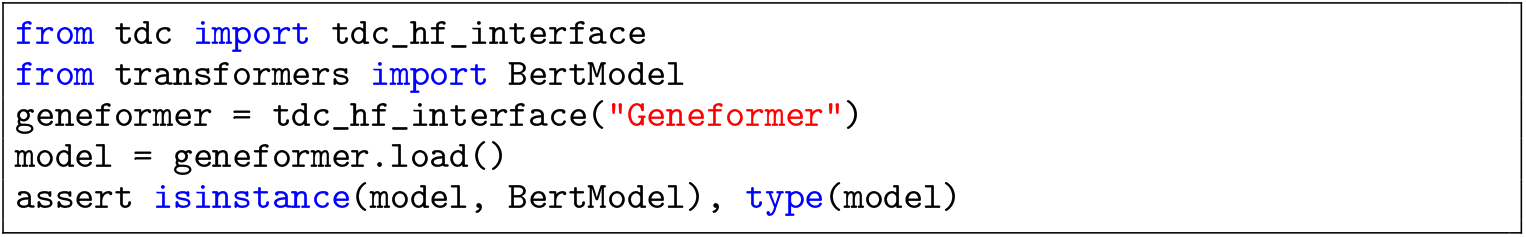
The below illustrates using the tdc model hub to download a foundation model [3]

**Listing 9:**
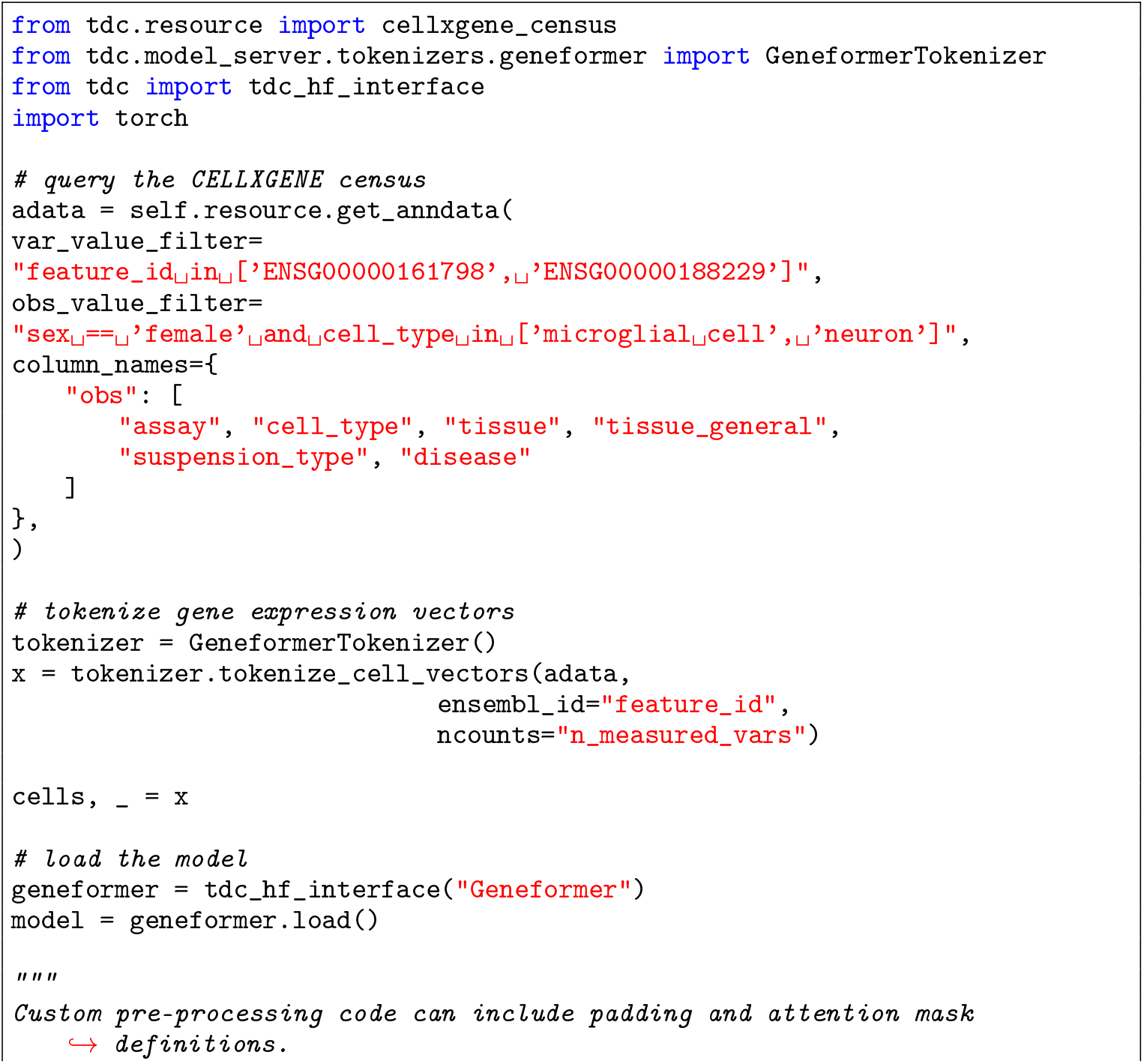
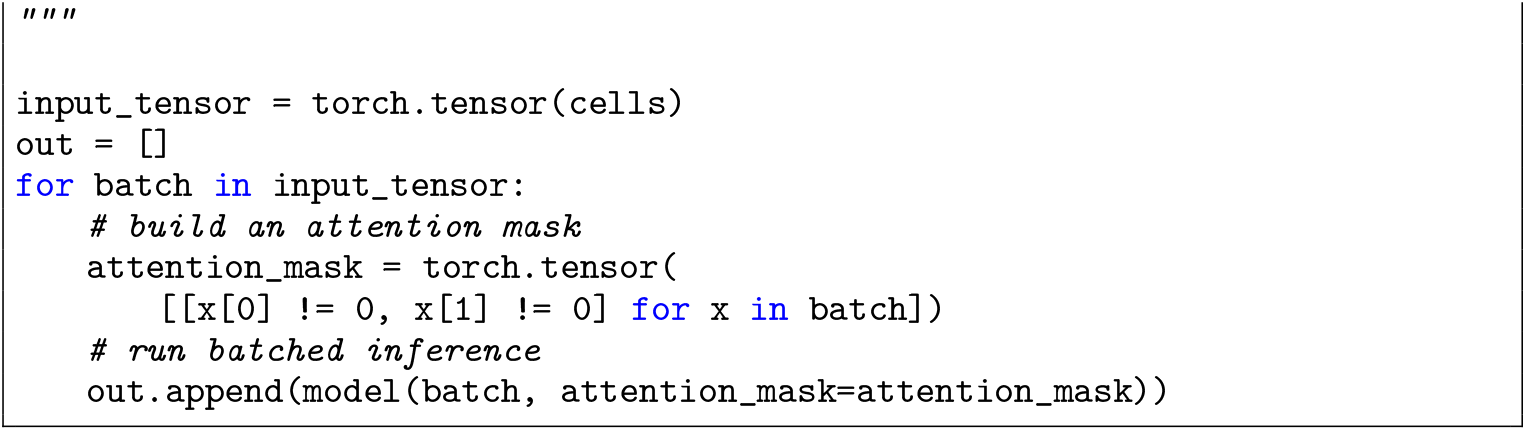
Beyond downloading a foundation model [3], the model server facilitates model inference across a range of datasets. Below an example integrating the TDC-2 CellXGene API with the model server.

#### 7.3.4 Running TDC-2 Benchmarks

We provide code for replicating all introduced benchmarks and testing other model performance on all TDC-2 tasks. We include here snippets for all introduced benchmarks.

**Listing 10:**
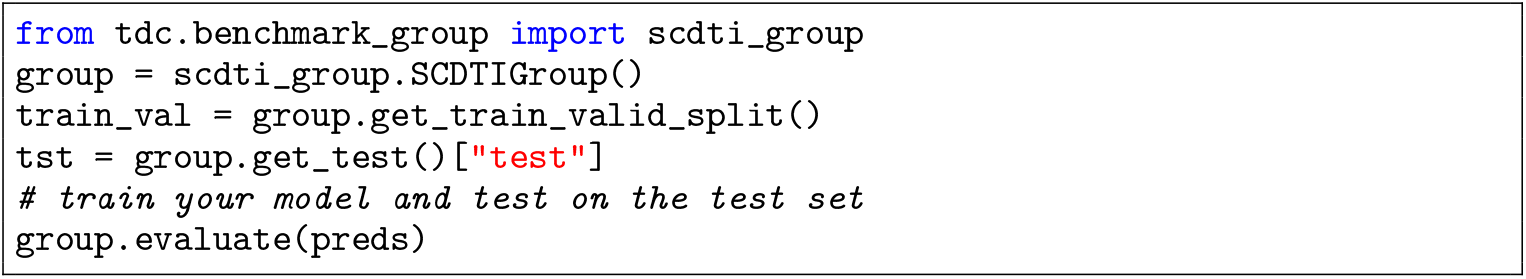
The below code illustrates how to retrieve the train, test, and val splits used for the TDC.scDTI benchmark

**Listing 11:**
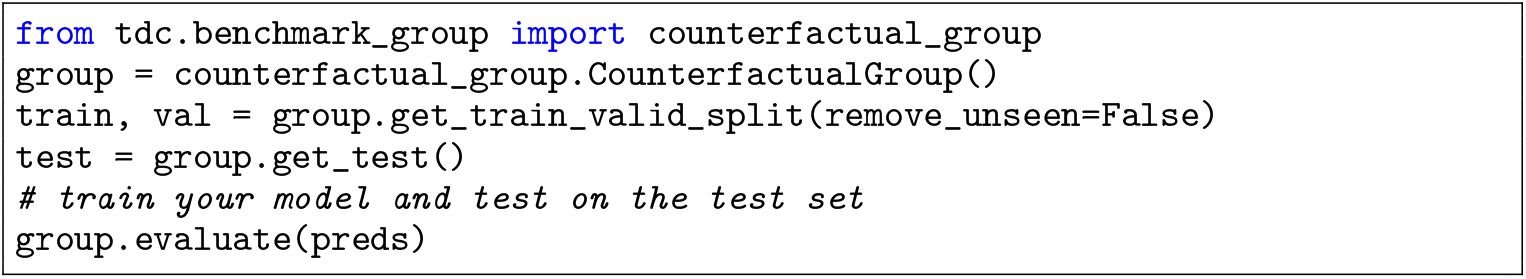
The below code illustrates how to retrieve the train, test, and val splits used for the TDC.PerturbOutcome chemical perturbation benchmark

**Listing 12:**
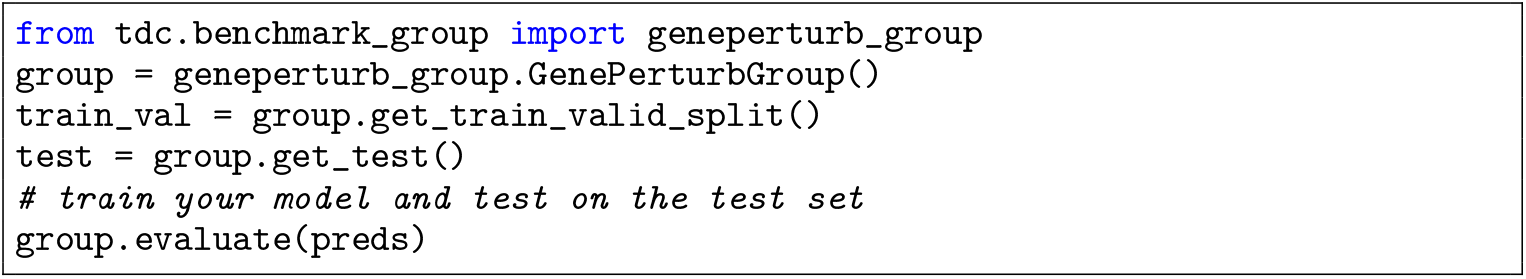
The below code illustrates how to retrieve the train, test, and val splits used for the TDC.PerturbOutcome genetic perturbation benchmark

**Listing 13:**
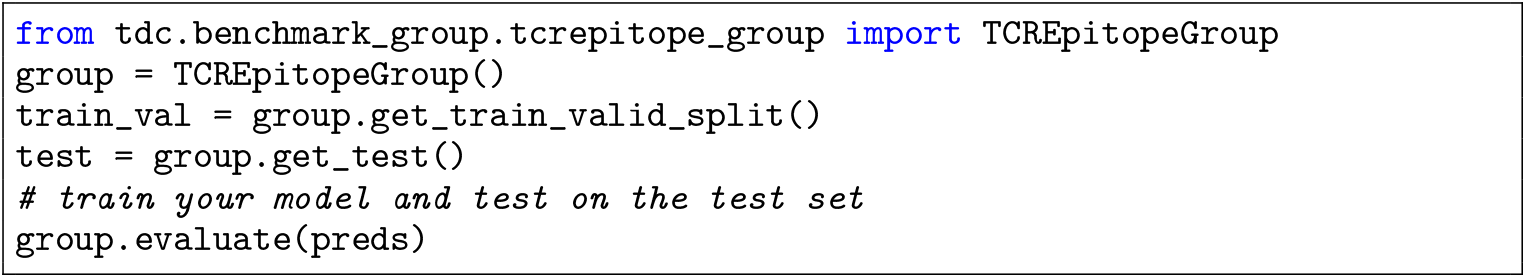
The below code illustrates how to retrieve the train, test, and val splits used for the TDC.TCREpitope benchmark

#### 7.3.5 External Links - Reproducibility

Here, we include pointers to external resources to reproduce the results reported in this manuscript.

##### Model Benchmarking

TDC-2 Benchmarking Tooling Code for Chemical Perturbations, https://github.com/mims-harvard/TDC/blob/main/tdc/benchmark_group/counterfactual_group.py

TDC-2 Benchmarking Tooling Code for CRISPR-based Perturbations, https://github.com/mims-harvard/TDC/blob/main/tdc/benchmark_group/geneperturb_group.py

Reproducing Benchmark Results for Clinical Trial Outcome Prediction, https://github.com/futianfan/clinical-trial-outcome-prediction

Evaluating Cell-Type-Specific Context Metrics for PINNACLE Across 10 Seeds, https://colab.research.google.com/drive/1gjZIfmF2Gmz3Nqm1uGP79l0AmsPAvj_5?usp=sharing

Evaluating Cell-Type-Specific Context Metrics for PINNACLE Across 10 Seeds. Outputs Referenced in PINNACLE [4] and its reproducibility documentation, https://drive.google.com/drive/folders/1QX05afMekucbtjl_07ZxZhgnKVH3OXMk?usp=sharing

Code for reproducing PINNACLE results [4], https://github.com/mims-harvard/PINNACLE/tree/main/evaluate

Reproducing TCR-Epitope results. Code for Benchmarking models in Section 3.3.1. A bash script for each negative sampling method is included for each TCR-Epitope model, https://drive.google.com/drive/folders/1O7G_h_06VDABM6U_Xt7otXPazK0XTAG9?usp=sharing

Reproducing Chemical Perturbation results. Code for Benchmarking models in Section 3.2.2 chemical perturbation section. A run_chemical_sc.py Python script is included for each model. Default settings were used from each model’s GitHub repository, https://drive.google.com/drive/folders/1RlBnRPmWFRQ6M_1EQ_FMwFb1Y8IjXoyC?usp=sharing

##### Leaderboards

TDC.PerturbOutcome Leaderboard, https://tdcommons.ai/benchmark/counterfactual_group/overview/

TDC.ProteinPeptide Leaderboard, https://tdcommons.ai/benchmark/proteinpeptide_group/overview/

TDC.scDTI Leaderboard, https://tdcommons.ai/benchmark/scdti_group/overview/

***Acknowledgments and Disclosure of Funding***

We thank Zitnik Lab, Pentelute Lab, and Kellis Lab members for their constructive input on the manuscript and contributions to TDC-2 datasets, benchmarks, and experiments.

We gratefully acknowledge the support of NIH R01-HD108794, NSF CAREER 2339524, US DoD FA8702-15-D-0001, awards from Harvard Data Science Initiative, Amazon Faculty Research, Google Research Scholar Program, AstraZeneca Research, Roche Alliance with Distinguished Scientists, Sanofi iDEA-iTECH Award, Pfizer Research, Chan Zuckerberg Initiative, John and Virginia Kaneb Fellowship award at Harvard Medical School, Aligning Science Across Parkinson’s (ASAP) Initiative, Biswas Computational Biology Initiative in partnership with the Milken Institute, Harvard Medical School Dean’s Innovation Awards for the Use of Artificial Intelligence, and Kempner Institute for the Study of Natural and Artificial Intelligence at Harvard University. M.M.L. is supported by T32HG002295 from the National Human Genome Research Institute and a National Science Foundation Graduate Research Fellowship. Any opinions, findings, conclusions or recommendations expressed in this material are those of the authors and do not necessarily reflect the views of the funders.

## Notes

https://tdcommons.ai/

https://github.com/mims-harvard/TDC

https://huggingface.co/tdc

### Competing Interest Statement

The authors have declared no competing interest.

### Summary of Updates

This version of the manuscript has been updated to match the existing accepted submission to NeurIPS 2024 Workshop on AI for new drug modalities.

